# Dominance is common in mammals and is associated with trans-acting gene expression and alternative splicing

**DOI:** 10.1101/2023.03.31.535109

**Authors:** Leilei Cui, Bin Yang, Shijun Xiao, Jun Gao, Amelie Baud, Delyth Graham, Martin McBride, Anna Dominiczak, Sebastian Schafer, Regina Lopez Aumatell, Carme Mont, Albert Fernandez Teruel, Norbert Hübner, Jonathan Flint, Richard Mott, Lusheng Huang

## Abstract

**Background:** Dominance and other non-additive genetic effects arise from the interaction between alleles, and historically these phenomena played a major role in quantitative genetics. However, today most genome-wide association studies (GWAS) assume alleles act additively.

**Methods:** We systematically investigated both dominance – here representing any non-additive effect - and additivity across 574 physiological and gene expression traits in three mammalian models: a Pig F2 Intercross, a Rat Heterogeneous Stock and a Mouse Heterogeneous Stock.

**Results:** In all species, and across all physiological traits, dominance accounts for about one quarter of the heritable variance. Hematological and immunological traits exhibit the highest dominance variance, possibly reflecting balancing selection in response to pathogens. Although most quantitative trait loci (QTLs) are detectable assuming additivity, we identified 154, 64 and 62 novel dominance QTLs in pigs, rats and mice respectively, that were undetectable as additive QTLs. Similarly, even though most cis-acting eQTLs are additive, we observed a large fraction of dominance variance in gene expression, and trans-acting eQTLs are enriched for dominance. Genes causal for dominance physiological QTLs are less likely to be physically linked to their QTLs but instead act via trans-acting dominance eQTLs. In addition, in HS rat transcriptomes, thousands of eQTLs associate with alternate transcripts and exhibit complex additive and dominant architectures, suggesting a mechanism for dominance.

**Conclusions:** Although heritability is predominantly additive, many mammalian genetic effects are dominant and likely arise through distinct mechanisms. It is therefore advantageous to consider both additive and dominance effects in GWAS to improve power and uncover causality.

## INTRODUCTION

Dominance arises from non-additive interactions between different alleles within a locus. The pathways that cause dominance still remain to be clarified, despite intense scrutiny [1–3]; suggested explanations include haplo-sufficiency (when a single working copy of one gene is sufficient for normal function) [4, 5], antimorphs (when the mutant product of one gene interacts and interferes with the normal product) [6–8], hypomorphs (when one allele has a partial or complete loss-of-function) [9] and antagonistic pleiotropy (when one allele is beneficial to some traits while deleterious to others) [10–12].

In the quantitative genetics model proposed by Fisher in 1916 [13–15], the genetic variance, or heritability, of a quantitative trait is partitioned into additive, dominant and epistatic components, each of which are aggregates of many smaller contributions within and between multiple causal loci. It follows that understanding dominance in the round depends on understanding dominance within each locus, and on clarifying the causal molecular pathways.

A given biallelic locus is additive if the phenotypic effect of the heterozygote is the mean of that of the two homozygotes. In this study, we define dominance to mean any non-additive within-locus interaction, classified as partial (PD), complete (CD) or over-dominance (OD), according to whether the phenotypic effect of the heterozygote lies within the range spanned by the homozygotes but is unequal to their average, or is equal to one of the homozygote effects, or is outside their range [16, 17]. Any possible additive or dominant relationship between a trait and a biallelic locus can be modeled by a combination of additive and complete dominance effects, and the presence of dominance in this wider non-additive sense is therefore testable by comparing the fit of a purely additive model to a model with both additive and dominance effects. Computational methods to detect dominance in GWAS have been developed by our group[18] and others[19–26].

Although most quantitative genetics studies assume additivity, dominance effects – where investigated - have been observed in genome-wide association studies (GWAS), heritability estimation, genomic selection and prediction. Crosses between strains often reveal the closely related phenomenon of heterosis. Dominance quantitative trait loci (QTLs) have been mapped in animals (cattle[27–33], pig[34–37], sheep[38], chicken[39, 40]), plants (maize[41–44], wheat[45], sunflower[46], *Arabidopsis*[47], *Primulina*[48]) and in a few studies in humans[49–53]. In cattle, where dominance effects have been investigated most intensively, recessive QTLs are known for lactation, growth and developmental traits[30, 32]. Similarly in pigs, dominance QTLs are associated with the number and weight of piglets born[37], number of teats[36], meat quality[35] and growth traits[34]. In plants, dominance QTLs are associated with disease resistance (shoot fly in maize [42] and stripe rust resistance of wheat[45]) and growth (leaf orientation in maize[44], flowering time in *Arabidopsis*[47] and sunflower hybrids[46], and hybrid male sterility of *Primulina*[48]). Consistent with these observations, a large fraction of dominance heritability is present across many traits in cattle (yearling weight[54], growth[55], milk production[56] and reproduction[57]) and pigs (sow longevity[58], daily gain[59, 60], number of teats[60], backfat[60, 61] and growth[61]).

In contrast with these findings, studies in humans have generally reported that both dominance variance components are small [62–65] and dominance associated loci [52, 53, 66] are relatively rare. One potential explanation might be the prevalence of low-frequency alleles in human and other large random mating populations. Crucially, experimental and artificially bred populations usually exhibit more limited haplotype diversity and higher allele frequencies. The power to detect dominance QTLs and to predict dominance phenotypes depends critically on the frequency of the rarer of the two homozygote genotypes and is consequently attenuated at lower allele frequencies, as shown in a recent study of recessive human disease [67].

There are also practical reasons why dominance is often ignored. First, modelling dominance requires extra degrees of freedom in fixed effects models, potentially reducing the power to detect purely additive effects. Second, if one is to model dominance effects using the mixed model framework, it follows that both additive and dominance variance components should be included in genetic background effects, which is computationally challenging. For these reasons, most GWAS in humans only consider additive variance components. Where dominance heritability is large this shortcut is potentially unsound.

Moreover, understanding the relationship between dominance at the level of gene expression and at the level of physiological phenotype may be key to establishing causal mechanisms. In yeast[68, 69], plants[70], flies[71], fish[72] and mice[73] dominance effects on gene expressions are associated with trans-acting control of gene expression and with structural variations such as translocations that silence or otherwise modulate the expression of genes[74]. In contrast, most cis-acting expression QTLs (eQTLs) are additive. These phenomena suggest how dominance arises at the molecular level, but their role in mammals is under-explored.

In this study, we systematically investigate dominance across physiological and gene expression traits in three mammalian species, namely pigs[75], rats[76] and mice)[77]. These populations were chosen because of the wealth of genotypes and phenotypes available combined with gene expression measured on large subsets of the same animals. Within each population, we analyse their dominance and additive genetic architectures through variance decomposition, QTL and eQTL mapping. Additionally, in the rats, we use RNA-seq data to relate dominance to the expression of alternative transcripts, revealing a novel mechanism for dominance.

## RESULTS

### 1. Population Characteristics

We integrated and analysed published multi-phenotype and multi-omics data from three mammalian stocks: (1) F2 intercross pigs (hereafter F2 pigs)[75], containing 1,005 progeny derived from 2 White Duroc boars mated with 17 Erhualian sows, with 253 complex traits measured related to growth, fatness, meat quality and blood. In addition, we analysed digital gene expression data of liver and muscle[78]; (2) Heterogeneous stock rats (HS rats)[76], encompassing 1,407 individuals descended from eight inbred founder strains (ACI/N, BN/SsN, BUF/N, F344/N, M520/N, MR/N, WKY/N and WN/N), in which 220 physiological traits and previously unpublished RNA-seq data of amygdala and heart samples were measured; (3) Heterogeneous stock mice[77] (HS mice), comprising 2,002 individuals descended from eight inbred founder strains (A/J, AKR/J, BALBc/J, CBA/J, C3H/HeJ, C57BL/6J, DBA/2J and LP/J), with measurements of 125 physiological traits and microarray gene expression data of hippocampus, liver and lung [79]. The chromosomes of the HS rats and mice are fine-grained mosaics of their respective inbred founder strains. The F2 pigs are not descended from inbred founders because individuals from the same pig breed are not genetically identical. The populations and datasets used, including the traits mapped in each population are summarized in Supplemental Tables S1 and S2.

Each phenotype was normalized, and the effects of covariates removed, as described previously[75–77]. All subsequent analyses used these normalized residuals. In each population, we removed SNPs with minor allele frequency (MAF) < 0.05, missing rate > 0.1 or if the rarest genotype occurred in fewer than 10 individuals. The numbers of SNPs passing these quality control steps was respectively 39,298 (pig), 244,786 (rat) and 9,142 (mouse).

### 2. Dominance accounts for about a quarter of genetic variance of organismal traits

We used these SNP sets to construct additive and dominance genetic relationship matrices (GRMs) and perform genetic mapping. We dissected the contributions of dominance to the 584 organismal traits measured across the three populations, by simultaneously estimating both additive ( *V_a_*) and dominance ( *V_d_*) genetic variance components from the GRMs in each trait using GCTA[62]. As each phenotype was scaled to have unit variance, these components also represent heritabilities (Supplementary Table S2). The relationship between *V_a_* and *V_d_* is shown as scatter plots (Fig. 1 a-c) and bar plots (Supplementary Fig. S1 a-c). Across 425 traits with nonzero dominance variance (*V_d_* > 0.05), *V_a_*/*V_d_* ≈ 3, i.e. dominance accounts for about one quarter of the genetic variance. In F2 pigs, *V̄_a_*: *V̄_d_* = 0.33: 0.11 across *n* = 163 traits. HS rats exhibit slightly higher average heritabilities (*V̄_a_*: *V̄_d_* = 0.48:0.24, *n* = 187), while in the HS mice they are lower (*V̄_a_*: *V̄_d_* = 0.22:0.08, *n* = 75).

**Fig. 1.**
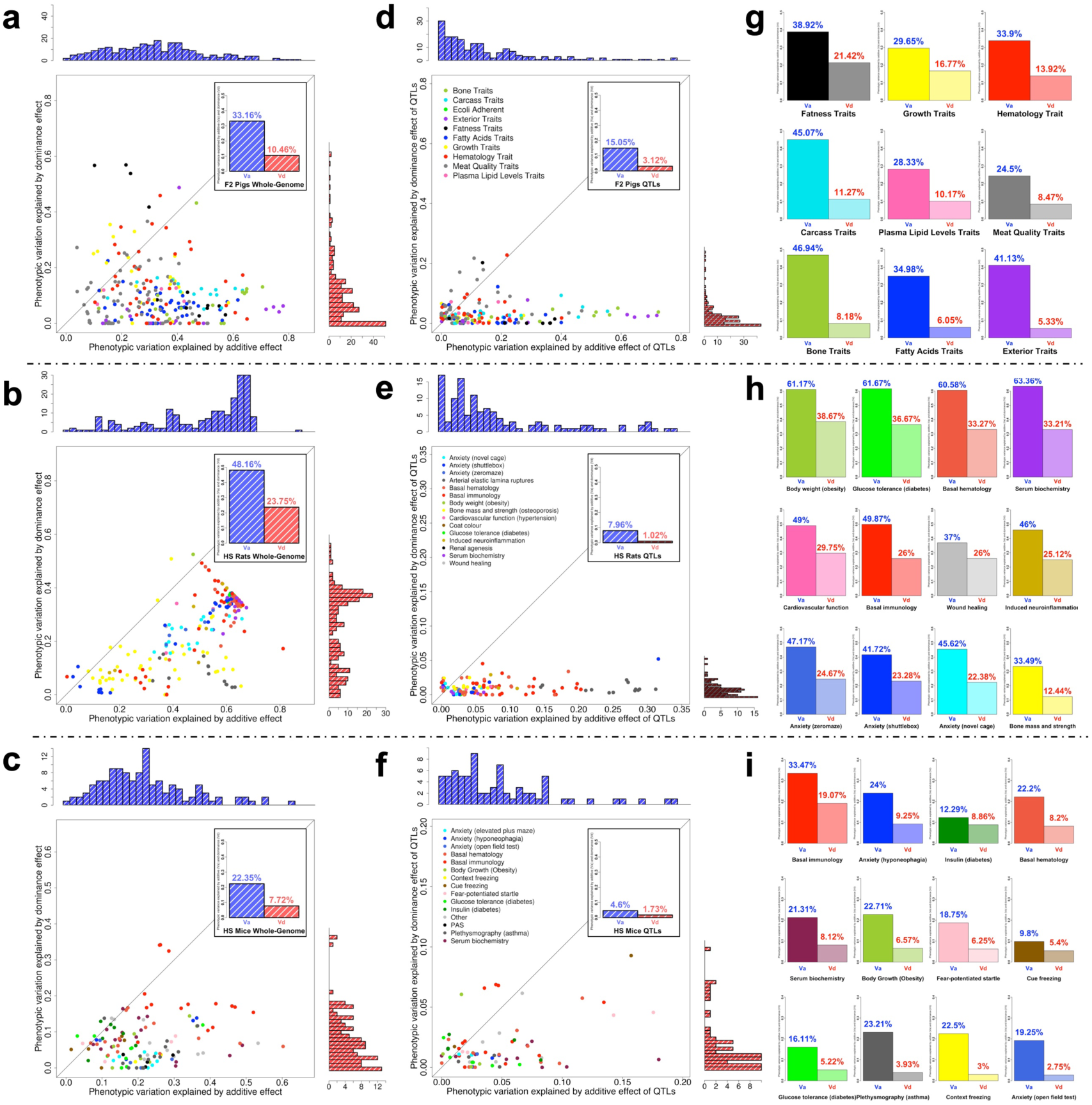
Comparison between additive and dominance heritabilities across multiple traits in three populations. (a-f) Additive Phenotypic variance component *V_a_* (x-axis) vs dominance component *V_d_* (y-axis), either estimated from genome-wide SNPs (a-c) or from accumulated significant QTLs (d-f) of 241 traits in F2 pigs, 206 traits in HS rats and 124 traits in HS mice respectively. Within each scatter plot, each dot denotes a trait, scaled to have unit variance so that variance components are also heritabilities 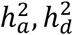. Dot colours show trait categories as tabulated in the left insets. The marginal histograms display distributions of *V_a_* (blue, top) and *V_d_* (red, right) of traits in each population, and their average values are indicated by upper-right inset bar plots. (g-i) Average *V_a_* (left) *V_d_* (right) for each population, colour-coded by trait category.

Thus, across all three populations and consistent with previous studies[75–77], additive genetic effects usually explain more phenotypic variance than dominance effects. However, many traits have important dominance contributions. In pigs, rats and mice, respectively, there are 52, 143, 15 traits (21.6%, 69.8%, 12.2%) where *V_d_* > 0.15, and 27, 8 and 13 (11.3%, 3.9%, 10.6%) where *V_d_* > *v_a_* (Supplementary Table S2, Supplementary Fig. S1). In F2 pigs, fatness, growth and hematology related traits exhibit the largest dominance effects (Fig. 1 a, g), whilst in rats and mice, immunology, hematology and serum biochemistry related traits have the strongest dominance effects (Fig. 1 b-c, h-j). As we show below, dominance and over-dominance QTLs are often detected in these traits.

### 3. Mapping dominance QTLs improves GWAS sensitivity

We used ADDO[18] to perform a mixed-model GWAS for each normalized trait, modelling both additive and dominance fixed effects at each focal SNP and including both additive, dominance and environmental variance components to model background effects. This Add-Dom (or AD) Model exhibits better calibration of *P*-values than a model solely including additive effects [18]. At each SNP, we compared the Add-Dom Model to either a null model with neither additive nor dominance effects or to a model with additive SNP effect only, but keeping the AD variance components, using ANOVA. We applied two Bonferroni *P*-value significance thresholds, (i) approximate 5% genome-wide significance (0.05/N_SNP_) (ii) suggestive significance (1/N_SNP_, corresponding to −log_10_ *P* = 4.5, 4.8, 3.9 in pigs, rats and mice respectively). To classify the effect type of each QTL, we first computed T-statistics *t_Add_* and *t_Dom_* from the Add-Dom Model’s additive and dominance parameter estimates (*β_Add_* and *β_Dom_*). Then, based on the base 2 logarithm of the absolute value of the ratio log_2_|*t_Dom_*⁄*t_Add_*|, we classified each QTL as (A-QTL) additive, (PD-QTL) partial-dominant, (CD-QTL) complete-dominant or (OD-QTL) over-dominant (Fig. S2.1 a-f). We used the log transformation log_2_|*t_Dom_*⁄*t_Add_*|, to visualise classifications (Fig. 2 a-c).

Using suggestive significance thresholds the AD Model detected 352 QTLs for 182 F2 pig traits, 179 QTLs for 119 HS rat traits and 116 QTLs for 73 HS mice traits (Fig. 2 d-f, T Supplementary able S4), of which 137, 87 and 31 QTLs were genome-wide significant, with average logP-thresholds of 5.8, 6.1 and 5.2 for pig, rat and mouse, respectively. We report results for the top SNP at each QTL in Supplementary Table S4. The log_2_|*t_Dom_*⁄*t_Add_*| ratios for each QTL, categorized by population and class of phenotype are plotted in Fig. 2 a-c, and the genome-wide distributions of QTLs shown as porcupine plots in Fig. S2.1 g-i. Similar proportions of each QTL type occur in each population, and dominance QTLs are common (Fig. 2 d-f, Supplementary Table S3). On average, 16% of suggestive QTLs are complete-dominant and 28.3% over-dominant. We annotated the peak SNPs of each QTL using the Variant Effect Predictor (VEP) tool[80], to predict consequences and related genes (Supplementary Table S4).

**Fig. 2.**
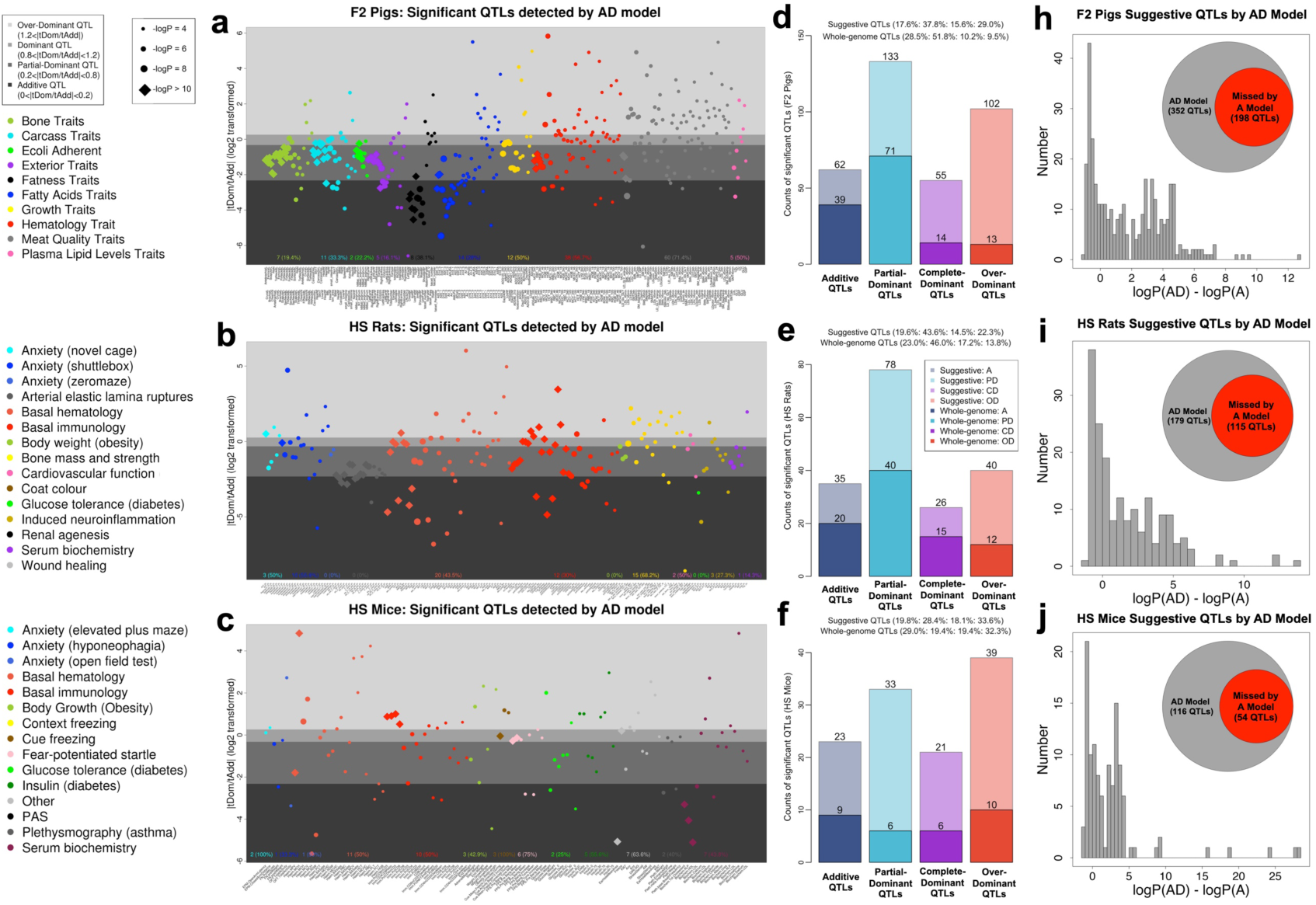
Classification and power improvement of dominance QTLs detected by the Add-Dom model. (a-c) Scatter plots showing QTLs detected at suggestive significant thresholds (one expected false positive per genome scan) identified in F2 pigs (a), HS rats (b) and HS mice (c). Each dot denotes a QTL, colors of dots represent trait categories, sizes of the dots represent the significance level (diamond points are those loci with -log_10_(*P*) values > 10) and vertical y-axis positions of the dots are their log_2_|*t_Dom_*/*t_Add_*| values, with background grey shades representing their classification from the bottom up as additive, partial-dominant, complete-dominant, and over-dominant. (d-f) Bar plots of the counts of additive (blue), partial-dominant (sky blue), dominant (purple) and over-dominant (red) QTLs in three populations. Light colors stand for the counts of suggestive significant QTLs and dark colors for whole genome significant QTLs. (h-j) Distribution of the difference between -log_10_(*P*) values of peak SNPs of suggestive significant QTLs detected by Add-Dom model compared to Add model in each population.

The AD Model has consistently greater power to detect QTLs (Supplementary Fig. S2.1 d-f) compared to the A Model (Supplementary Fig. S2.1 a-c), especially for CD and OD QTLs. Among all suggestive QTLs, 44.3% are detected by the AD Model but absent from the A Model (Fig. 2 h-j, 43.8%, 35.8%, 53.4% for pig, rat and mouse). These comprise 100 (98%) OD QTLs and 39 (70.9%) CD QTLs in F2 pigs, with similar counts of 15 (57.7%) and 35 (87.5%) in HS rats, and 16 (76.2%) and 34 (87.2%) in HS mice. In addition, the AD Model improved the -log_10_(*P*) values of 119 (33.8%) pig QTLs by more than 4 units compared to the A model, and similarly in for 52 (29.1%) rat QTLs and 40 (34.5%) mouse QTLs (Table S4). Most newly detected or improved QTLs relate to hematology and immunology traits, consistent with the variance decomposition results.

We also investigated the power of the “D Model” which mapped purely dominant QTLs (Methods). Among suggestive SNPs detected by the AD, A or D Models (Supplementary Fig. S2.2 a-c) there was a large increase in -log_10_(*P*) values of significant SNPs (either uniquely or concurrently) detected by AD compared to both A (Supplementary Fig. S2.2 d-f) and D Models (Supplementary Fig. S2.2 g-i). Thus the AD model is uniformly more powerful than either simpler model.

Fig. 3 shows six representative QTLs that are improved or only detected by AD, for F2 pigs (Fig. 3 a-b), HS rats (Fig. 3 c-d) and HS mice (Fig. 3 e-f). Further examples for F2 Pigs are shown in Supplementary Fig. S3.1, (meat quality traits), and for HS rats in Supplementary Fig. S3.2 (immune cell traits), and for HS mice in Fig. S3.3 (immune cell traits). In the latter case, two dominance QTLs related to mouse T cell traits each localize to a potential causal gene *Bat3*, which is over 20 logP units more significant than the QTL found by the A-model in the neighboring gene *Myo1f*, thereby showing the AD model improves the detection and resolution of QTLs.

**Fig. 3.**
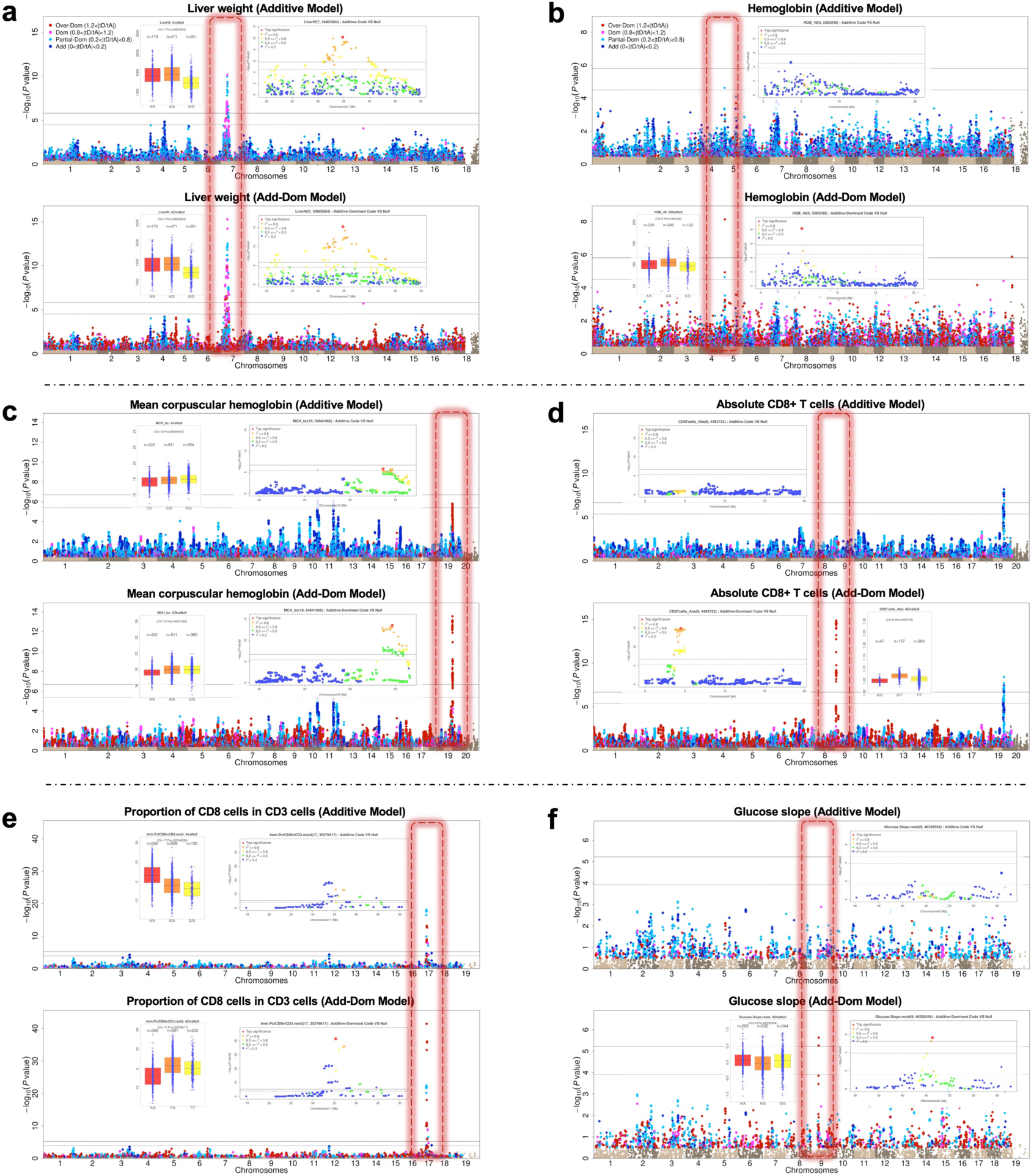
Novel and Enhanced dominance QTLs. (a, b): body weight and hemoglobin in F2 pigs. (c, d): mean corpuscular hemoglobin and absolute CD8+ T cells in HS rats. (e, f): proportion of CD3+ cells expressing CD8+ and glucose slope in HS mice. Within each part (a-f), the upper and lower panels show Manhattan plots for Add vs Add-Dom model respectively. Each panel includes insets representing the regional QTL plot and the phenotypic distribution of the peak SNP. Within each Manhattan plot, the QTL is marked by a red dotted rectangular frame, left column (a, c, e) for enhanced eQTLs and right column (b, d, f) for novel eQTLs. SNPs with -log_10_(*P*) > 0.5 are colour-coded by dominance (Blue: additive, Sky blue: partial-dominant, Purple: complete-dominant, Red: over-dominant). Color coding in regional Manhattan plots is different and represent linkage disequilibrium (r^2^) with the top SNP.

For each trait, we aggregated the variance explained by the peak SNPs at all independent QTLs, partitioned into additive (*V_a_QTL_*) and dominance (*V_d_QTL_*) contributions (Fig. 1 d-f), and compared them with the genome-wide variance components computed using GCTA (Fig. 1 a-c). Each population showed the expected missing heritability, where less variance was explained by QTLs.

We identified pleiotropic dominance QTL hotspots (18 in pig, 9 in rat and 9 in mouse; Supplementary Table S5-1 to S5-3). In F2 pigs, chr7: 34.8Mb-35.1Mb is associated with many growth-related traits (e.g. ear weight, bone length, skin thickness and carcass length), and the over-dominant hotspot chr11: 16.3Mb-66.8Mb is associated with many pork color traits, serum glucose level and hematocrit. Similarly, in HS rats, the hotspot at chr9: 3.78Mb-4.67Mb - distinct from the rat major histocompatibility complex (MHC) locus - is associated with immunology traits. In HS mice, a hotspot in chr9: 72Mb-111Mb is associated with hematology traits while the MHC hotspot chr17: 33.7Mb-37.2Mb is associated with immunology traits.

Among traits measured on more than one species, there are six groups of potentially homologous QTLs across all three species (Supplementary Table S6), namely (i) CD4/CD8 T cells of rat and mouse, (ii) heart weight of pig and rat, (iii) body growth or body weight of three species, (iv) glucose tolerance of rat and mouse, (v) various serum biochemistry related traits of rat and mouse (e.g. HDL, LDL, Cholesterol, Urea and so on) and (vi) hematology traits in all three species (e.g. HCT, HGB, MCH, MCHC, MCV, PCT, RDW and so on). The most significant examples are for MCV (Supplementary Fig. S4.1) and MCH (Supplementary Fig. S4.2). Many other examples replicate between two species (Supplementary Fig. S4.3). For example, within the syntenic rat and mouse MHC regions we observe dominance for many immunological traits. In pigs dominance loci do not appear to be syntenic with those in rodents, but no F2 pig immune system traits were available for analysis.

### 4. Gene expression is strongly influenced by dominance effects

We next investigated the impact of dominance on gene expression. We evaluated seven tissues across the three populations: F2 pigs (liver and muscle), HS rats (amygdala and heart) and HS mice (hippocampus, liver and lung), via variance decomposition with GCTA and eQTL mapping with ADDO. For rat RNAseq data we made separate analyses for gene and transcript expression, where a gene’s expression aggregates those of its transcripts.

The heritability of most expression traits was lower than for physiological traits, but surprisingly a larger fraction of the variance was accounted for by dominance, even though, as we describe below, most cis-eQTLs are additive. Figure 4a shows the averaged relative proportions of gene expression variances across species and tissues in comparison with the physiological phenotypes shown in Figure 1d-f. In pigs, additive variance components *V_a_* were generally larger than dominance components *V_d_*. Interestingly the reverse is the case in rats and mice. Although the standard errors of the estimated variance components are large, t-tests showed the average differences between *V_a_* and *V_d_* were significant (*P*-value < 10^−4^).

**Fig. 4.**
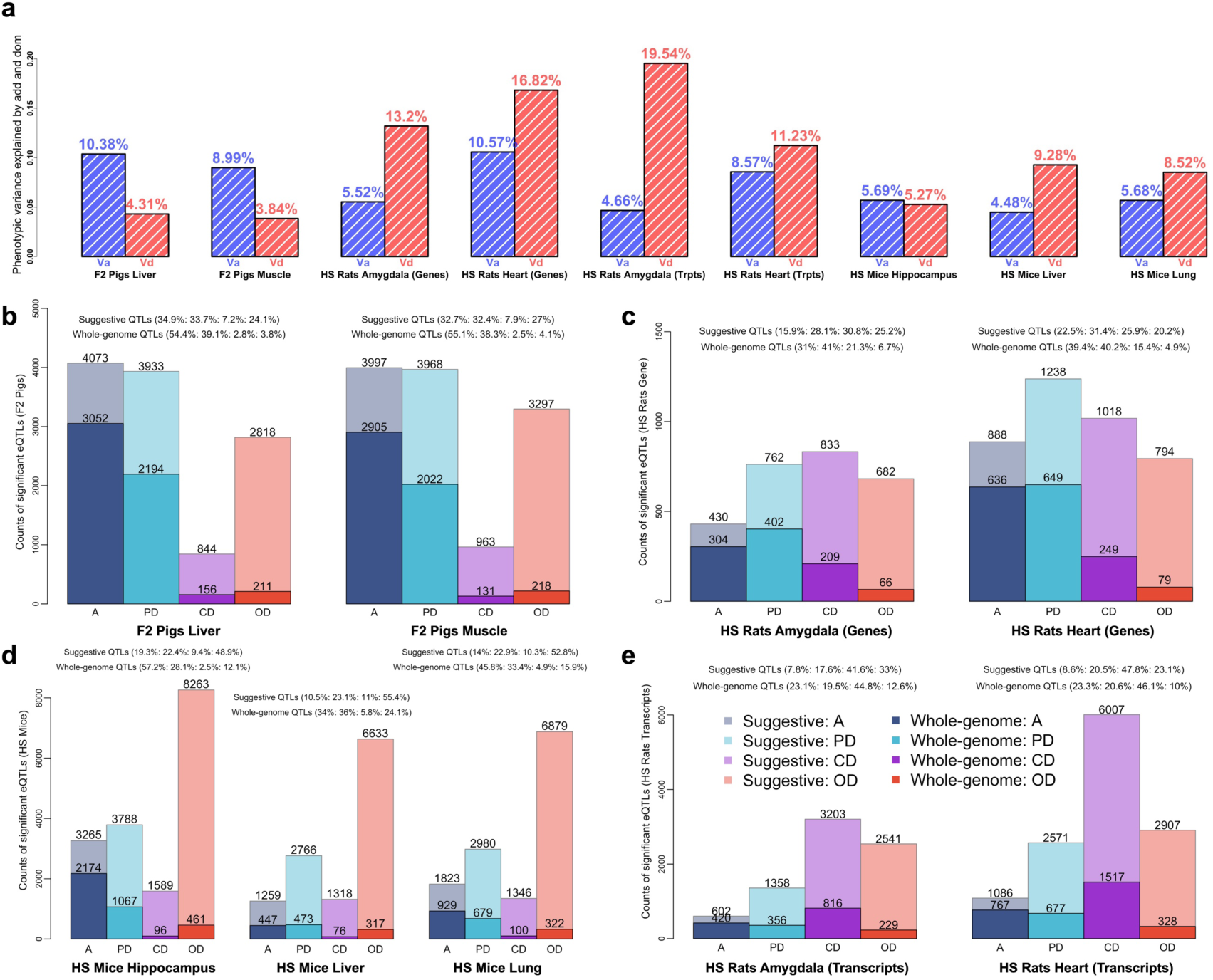
Dominance effects in gene expression. (a) Bar charts show the average heritabilities of gene expression variation for additive (*V_a_*:blue) and dominance (*V_d_*:red) effects per species and tissue. (b-e) Bar charts of the proportions of additive (blue), partial-dominant (sky blue), dominant (purple) and over-dominant (red) eQTLs in different tissues in (b) F2 pigs (liver and muscle), (c) HS rats (genes expression of amygdala and heart), (e) HS rats (transcripts expression of amygdala and heart) and (d) HS mice (hippocampus, liver and lung. Light shades: counts of suggestive significant eQTLs. Dark shades: whole genome significant eQTLs.

To eliminate potential inaccuracies when variance components are very small, we repeated the analysis restricted to genes where both *V_a_* > 0.05 and *V_d_* > 0.05. We observed *V_d_* > *V_a_* in 33.6% pig genes, 67.8% rat genes and 56.3% mouse genes (Supplementary Table S7). Many of these genes exhibit very high dominance (defined as *V_d_* > 0.15), including (1) 405 liver and 295 muscle genes in pig; (2) 1,651 amygdala and 9,491 heart genes in rat; (3) 887 hippocampus, 2,540 liver and 2,482 lung genes in mouse.

We then mapped eQTLs using the same methodology as for physiological traits (Supplementary Table S9-1 to S9-9), using thresholds ranging from lax (-log_10_(*P*) = 5.8, 6.3, 5.1 for pig, rat and mouse) to stringent (-log_10_(*P*) =8.5). We mapped thousands of eQTLs in each tissue, as detailed in (Supplementary Table S8-1 and Fig. 4 b-e). We applied a series of thresholds of increasing stringency in order to understand how the incidence of dominance eQTLs varied between cis and trans eQTLs at different thresholds. Supplementary Table S8-2 gives the proportions of novel eQTLs (i.e. only detectable by the AD model) at different thresholds. Notably, many trans-eQTLs were novel; at suggestive thresholds, 42.3%, 81.1% and 71.1% trans-eQTLs are novel in pig, rat and mouse. At more stringent thresholds - and therefore among progressively fewer eQTLs - we still observed high fractions of novel trans-eQTLs (5.7%, 61.5% and 44.7% respectively).

In contrast with our heritability results, at stringent thresholds most eQTLs are additive or partial dominant. However, 66.8% of pig eQTLs are A&PD-eQTL, over half of rat eQTLs are CD&OD (51.1% and 72.8% for gene and transcript eQTLs respectively), and 52.4% of suggestive mouse eQTLs are OD.

Consistent with other studies, we mapped numerous trans eQTL hotspots which are also strongly enriched for dominance effects (Supplementary Table S10-1 to S10-9). At threshold -log_10_(*P*) > 10, there are 537 and 479 hotspots affecting 1,919 liver genes and 1,721 muscle genes of F2 pigs; (2) 67, 155, 57 and 127 hotspots 196 amygdala genes, 456 heart genes, 160 amygdala transcripts and 368 heart transcripts of HS rats; (3) 400, 71 and 168 hotspots affecting 1,674 hippocampus genes, 224 liver genes and 579 lung genes of HS mice. Figure 5 shows a dominance gene-level hotspot at chr10: 85Mb-86Mb in HS rat heart. This hotspot has two cis-eQTLs for the transcription factors (TF) *Tbx21* (over-dominant) and (e) *Nf32l1* (partial-dominant), that link to six trans-eQTLs. Scatter plots of the expression of the trans and cis-eQTLs reveals a complex pattern of correlations suggesting each cis-eQTL is associated with distinct trans-eQTLs (*Klrb1*, *Znf683*, *Cdh17* with *Tbx21, Gzmb*, *Cd160*, *F1lnm2* with *Nf32l1*), Fig. 5b-h. Correlation analysis of these transcripts shows a potential regulatory relationship. Supplementary Figure S5.1 shows the corresponding transcript-level hotspot. Supplementary Figures S5.3-S5.6 show further examples.

**Fig. 5.**
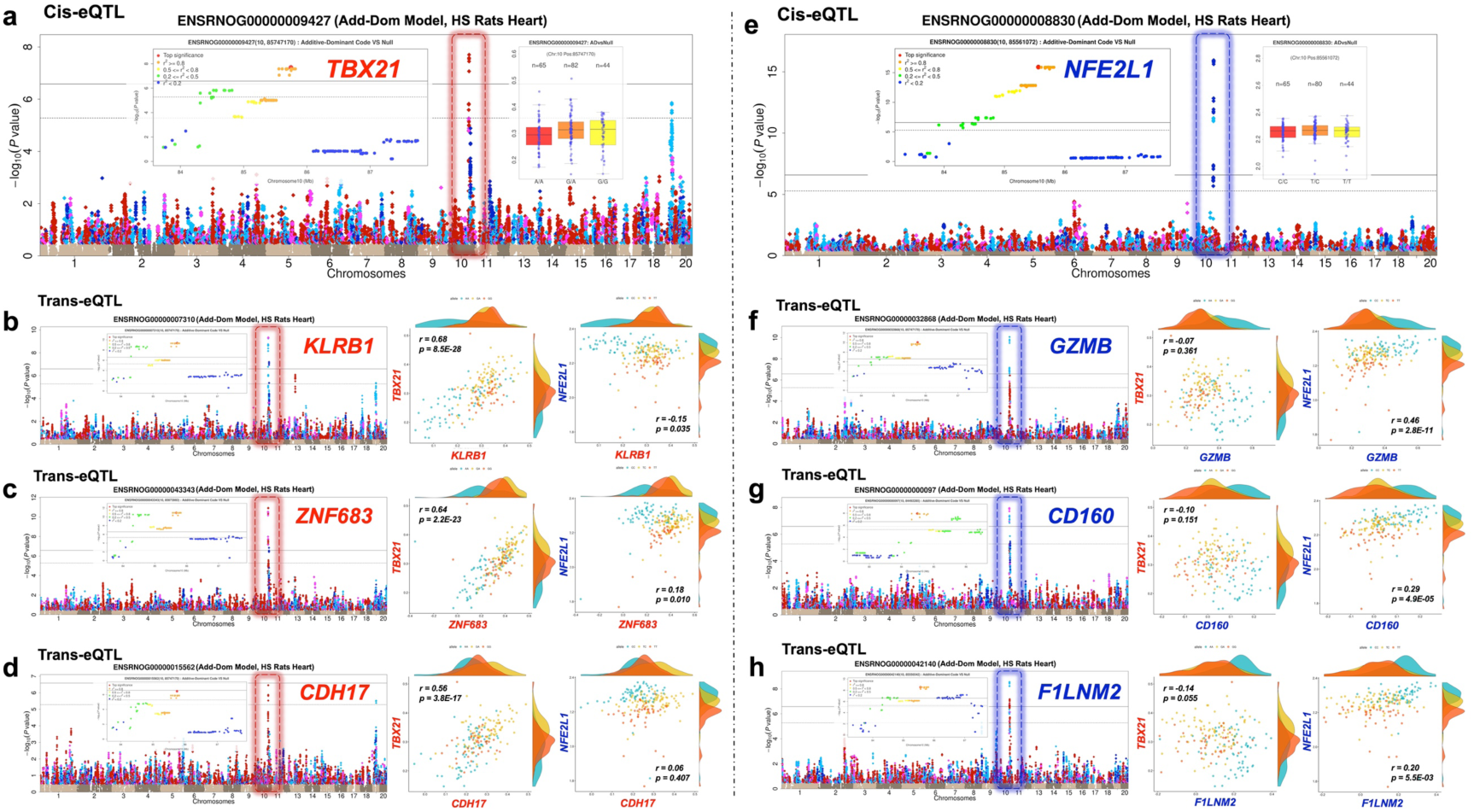
Dominant cis and trans eQTLs at the hotspot chr10:85Mb-86Mb in HS rat heart. (a, e) Two cis-eQTLs with different inheritance modes, (a) *Tbx21* (over-dominant) and (e) *Nf32l1* (partial-dominant). (b-d) Manhattan plots of over-dominant trans-eQTLs for genes *Klrb1*, *Znf683*, *Cdh17*. (f-h) Manhattan plots of partial-dominant trans-eQTLs for genes *Gzmb*, *Cd160*, *F1lnm2*. Within each Manhattan plot, the eQTL is marked by a dotted rectangular frame, with the same colour as the peak SNP dot (Blue - additive; Sky blue - partial-dominant; Purple - complete-dominant; Red - over-dominant), and all linked SNPs with -log10(P) > 0.5 are coloured the same. The regional Manhattan plots of the peak signal of each eQTL and the scatter plots of two cis-eQTLs are also shown as insets. The pairs of scatter plots to the right of each Manhattan plot compare the expression of each trans-eQTL with *Tbx21, Nf32l1.* Each dot represents one animal, colour-coded by the genotype of the peak SNP.

Hundreds of co-localised dominance eQTLs regulate the same gene in different tissues, (tc-eQTL), as distinct from tissue-specific eQTLs that regulate different genes in different tissues, (ts-eQTL). These are tabulated in Supplementary Table S11-1 to S11-6 using the stringent threshold -log_10_(*P*) > 10. In pigs there are 765 tc-eQTLs and 6,109 ts-eQTLs between liver and muscle genes. In rats between amygdala and heart genes there are 131 tc-eQTLs and 621 ts-eQTLs, and 92 tc-eQTLs and 487 ts-eQTLs in the corresponding transcripts. In mice there are 167, 411, 167 tc-eQTLs and 1273, 2751, 652 ts-eQTLs in hippocampus-liver, hippocampus-lung, liver-lung comparisons respectively. Representative examples of tc-eQTLs and ts-eQTLs are shown in Supplementary Fig. S5.7. They include: (Supplementary Fig. S5.7 a-b) an OD tc-eQTL for OD expression of *GP1* in liver and muscle in F2 pigs; (Supplementary Fig. S5.7 i-j) a PD tc-eQTL for *H2-Q1* expression levels of hippocampus and liver in HS mice; (Supplementary Fig. S5.7 m-n) a PD-eQTL for *9030612M13Rik* expression levels of liver and lung in HS mice.

### 5. Trans-acting enrichment among dominance eQTLs

We cross-tabulated additivity vs dominance against trans vs cis eQTLs using Fisher exact tests. To simplify results, we grouped A and PD-eQTLs into “generalized additive eQTLs” (G-Add eQTL) and CD and OD-eQTLs into “generalized dominance eQTLs” (G-Dom eQTL). Our results are summarized in Supplementary Table S12-1 and S12-2. Overall, and consistent with other studies, most cis-eQTLs are additive whilst trans-eQTLs are enriched for dominance effects, although it is not the case that most trans-eQTLs are dominant. Across a range of thresholds (from 5.5 to 8.5), there are statistically significant enrichments (all the *P*-values across seven tissues are smaller than 0.0001 when the significance thresholds = 5.5) for trans-acting effects among G-Dom eQTLs (Supplementary Table S12-1), and for cis-acting effects among G-Add eQTLs.

The spatial distributions of transcript-level HS rat eQTLs are shown in Figure 6, and similar results for gene-level eQTLs in rats, pigs and mice are in Supplementary Figures S6.1-S6.3. Figure 6a-j shows the positions of eQTLs SNPs vs their associated transcripts, filtered by dominance type in HS rat amygdala and heart. There is a strong diagonal band of G-Add cis-eQTLs whereas G-Dom eQTLs are evenly distributed, not-withstanding the presence of several vertical “trans-hubs”. The phenomenon was most noticeable using less stringent thresholds (-log_10_(*P*) = 4.7) where counts are large, but it persisted at more stringent thresholds up to -log_10_(*P*) value of 8.5 (Fig. 6k, Supplementary Table S12). Overall, at suggestive significant transcript levels in amygdala and heart respectively, 94.8% and 94.7% cis-eQTLs are G-Add. In contrast, 99.3% and 99.2% dominance eQTLs are trans-acting.

**Fig. 6.**
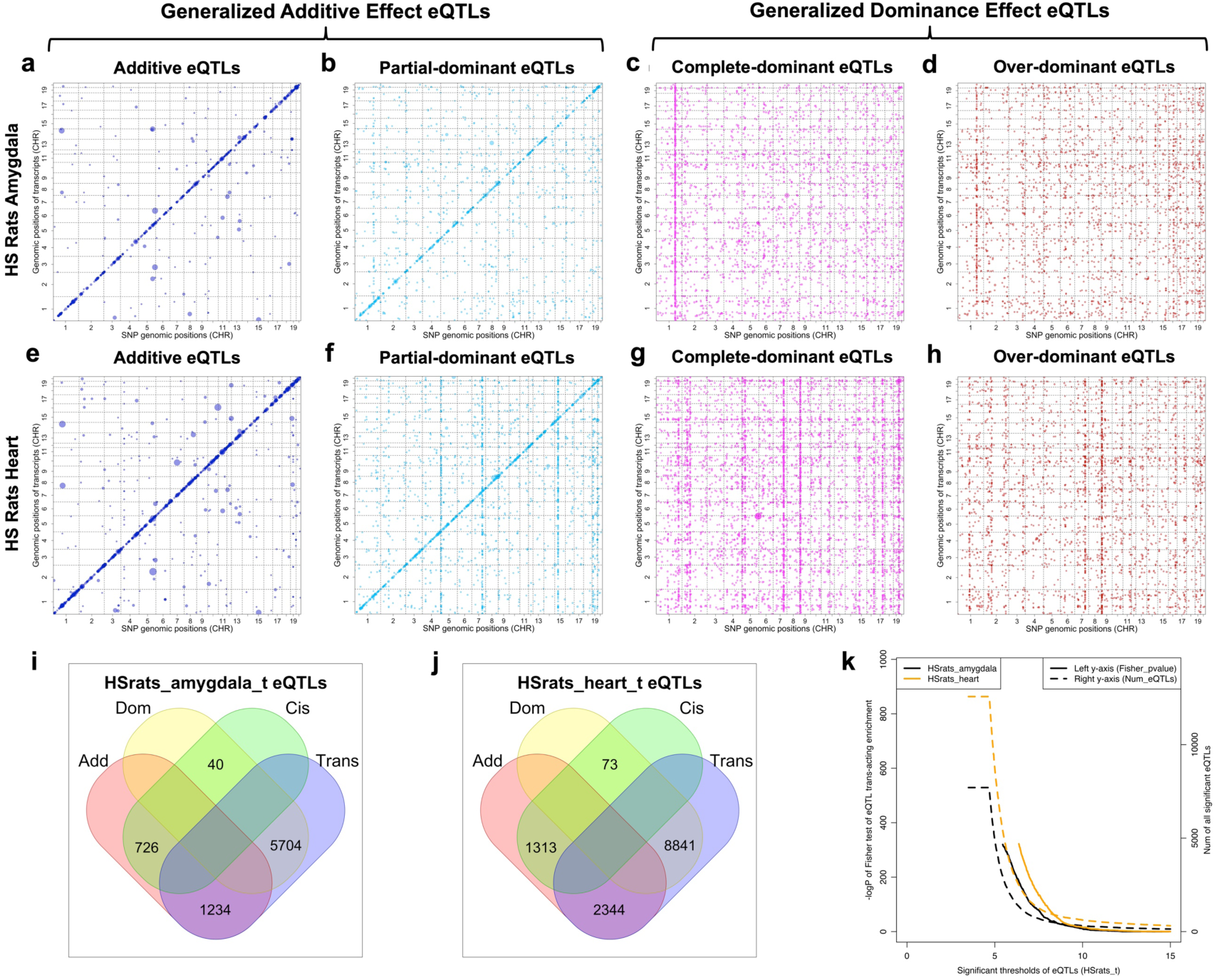
Trans-acting enrichment among dominant eQTLs of amygdala and heart tissues in HS rats. (a-h) eQTL locations of transcript level eQTLs, filtered by dominance type. Each dot represents an eQTL significant at suggestive level (i.e. one false positive expected per transcript). x-axis: eQTL chromosome positions (bp), y-axis: physical gene location. First row (a-d): amygdala; Second row (e-h): heart. The four columns represent dominance types (blue: additive A, sky blue: partial-dominant PD, purple: complete-dominant CD, red: over-dominant OD). (i: amygdala, j: heart) Venn diagrams comparing eQTL cross-classifications, (1) Generalised Additive eQTLs, (A-eQTLs and PD-eQTLs), (2) Generalised Dominance eQTLs, (CD-eQTLs and OD-eQTLs) vs (3) Cis-acting eQTLs, (4) Trans-acting eQTLs. (k) Dominance trans-acting enrichment (left y-axis, solid lines) and the counts of significant eQTLs (right y-axis, dashed lines) under different -log_10_(P) eQTL significance thresholds (x-axis). Enrichment is quantified by the -log_10_(P) values of Fisher exact test between dominance (Add/Dom eQTLs) and regulation types (cis/trans-acting) of significant eQTLs within rat amygdala (black) and heart (orange), respectively.

### 6. Dominance enrichment among genes with multiple transcripts

HS rat gene expression was measured by RNA-seq, which made it possible to distinguish expression levels of alternative transcripts, and to investigate the dominance enrichment of transcript-based eQTLs. Using the Rn4 reference annotations of 34,721 transcripts in 24,688 genes, 74.5% of genes express only one known transcript, while 22.2% (5,489 genes) express two or three transcripts (Table S13-2). In amygdala, we detected a higher proportion of G-Dom eQTLs among genes with multiple transcripts (ratio of G-Dom: G-Add eQTLs = 3.23, Supplementary Table S13-1) compared to genes with only one transcript (G-Dom: G-Add = 2.16, Supplementary Table S13-1). Chi-squared tests of enrichment were significant in both amygdala (*P*-value = 7.9*10^−11^) and heart (*P*-value = 5.1*10^−3^). Supplementary Table S13-3 to S13-8 list the QTL positions for each gene with their corresponding transcripts, including 521 amygdala and 828 heart gene-based eQTLs with two or three transcripts. Supplementary Tables S13-9 and S13-10 show examples of antagonistic and synergistic transcript expression in amygdala and heart.

Different transcripts of the same gene are most often associated with different SNPs; across the 4,086 rat genes with exactly two transcripts, 1257 amygdala and 1785 heart genes contain transcript-based eQTLs, but only 34 amygdala and 94 heart genes share eQTLs for both transcripts. The relative expression of different transcripts for the same gene can be either antagonistic (Fig 7 and Supplementary Fig. S7) or synergistic (Supplementary Fig. S7.2). Remarkably 3,942 amygdala and 5,659 heart genes have no gene-level eQTL but exhibit transcript level eQTLs, and this phenomenon is even more noticeable in genes with more transcripts; across gene sets with 1/2/3 transcripts we observed respectively 13.1%, 25.9%, 33.6% cases in amygdala and 20.2%, 34.1% and 39.8% in heart. G-Dom eQTLs explain 82.6% and 78.8% of these cases in amygdala and heart respectively. Overall, we observed statistically significant enrichments of dominance effects in genes with multiple transcripts (P=7.9*10^−11^ in amygdala and P=0.0051 in heart; Supplementary Table S13-1), thereby suggesting a mechanism by which dominance gene expression can arise.

**Fig. 7.**
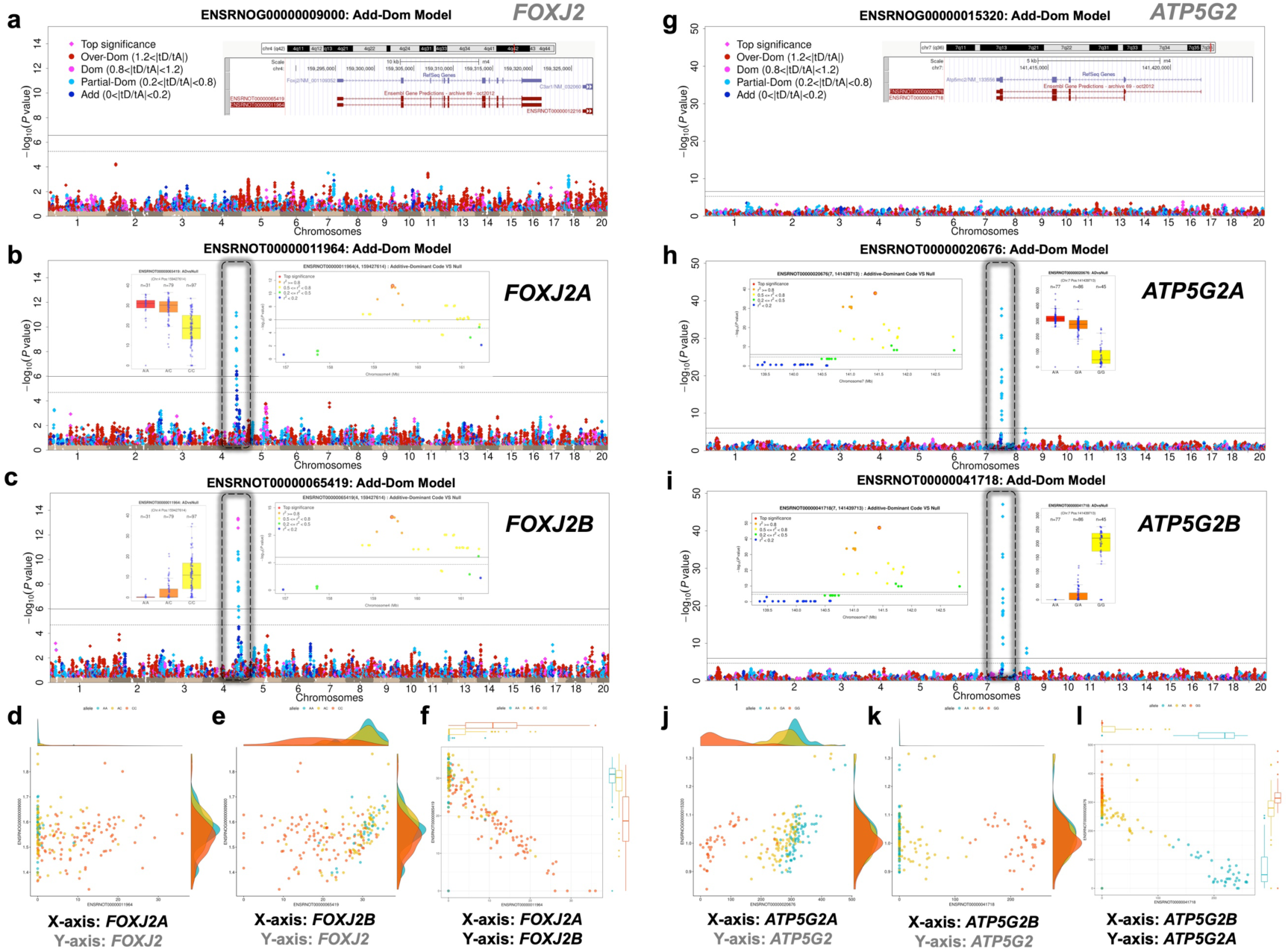
Transcript-specific antagonistic dominant eQTLs. (a-c, g-i) Manhattan plots of *Foxj2* and *Atp5g2*, based on their overall gene expression level in HS rat amygdala, and showing no genome-wide significant eQTLs; (a): *Foxj2* (g): *Atp5g2,* with their associated transcripts’ expression levels (b): *Foxj2A,* (c): *Foxj2B,* (h): *Atp5g2A* (i): *Atp5g2B* respectively, and showing cis eQTLs. Plot layouts are as for Fig 3, showing Manhattan plots colour coded by dominance type, regional QTL plots and phenotype-genotype distribution at peak SNPs. The transcript structures for *Foxj2*, *Atp5g2* from UCSC Genome Browser are inset. (d-f) Scatter plots of the correlations of expression levels between *Foxj2A vs. Foxj2* (d), *Foxj2B vs. Foxj2* (e) and *Foxj2A vs. Foxj2B* (f). (j-l) Scatter plots of the correlations of expression levels between *Atp5g2A vs. Atp5g2* (j), *Atp5g2B vs. Atp5g2* (k) and *Atp5g2A vs. Atp5g2B* (l). Within each scatter plot, one dot represents one sample, the dot colours indicate the genotype at the corresponding peak SNPs.

We show examples from rat amygdala of antagonistic transcript eQTLs (*Foxj2*, Fig. 7 b-c; *Atp5g2*, Fig 7 h-i). In both cases, alternate transcript levels are compensatory – and regulated by dominant variants - such that overall gene expression (Fig. 7 a, g) is not under genetic control whilst each transcript is under strong control. Scatter plots of corresponding transcript levels (colour-coded by genotype) illustrate the antagonistic effects of a SNP on different transcripts from the same gene (Fig. 7 d-f, j-l). Supplementary Figure S7-1 (a-d) presents two more antagonistic examples for *Rpl14* and *LFI44* verified in both amygdala and heart, which has no gene-level eQTL but an antagonistic pair of transcripts with a common eQTL. Four examples of synergistic transcript pairs are shown in Supplementary Figure S7-2 (a-d), including *Crot* and *Slc39a12* in amygdala and *Sppl2a* and *Rt1-m6-2* in heart, where alternate transcripts are controlled independently and involving additive effects.

## DISCUSSION

We have demonstrated here that understanding the prevalence, causes and consequences of genetic dominance not only clarifies the genetic architecture of complex traits[47, 51, 81] but also improves the power to detect associations. Our work complements earlier studies that have improved genomic prediction in theoretical simulation studies[82, 83] and in practical breeding applications for animals[84, 85] and plants[86, 87].

In summary, about a quarter of the heritability of the physiological traits we analysed is attributable to dominance, and there are non-additive effects of various types at over half of the QTLs we mapped. The fractions of QTLs and eQTLs showing dominance effects are similar across species. We observed an enrichment of trans-acting effects in generalized dominance eQTLs, which has been previously reported in yeast[68, 69], plant[70], fly[71], fish[72] and mouse[73]. We also observed an enrichment for dominance in genes that expressed multiple transcripts, which to our knowledge is a novel finding.

There are three specific advantages to model both additive and dominance genetic effects as we have done. First, the strategy detects more genetic associations (Supplementary Fig S2.1-S2.2). We mapped consistently more (44.3%) associations than additive modelling alone, despite the burden of fitting additional parameters. Most of these novel QTLs are completely (68%) or over (90.9%) dominant (Fig. 2). This effect is greatest in immunological and hematological traits, where some QTLs have negligible additive signal. Many Mendelian blood disorders in humans are dominant or recessive[88]. Although dominance studies of human traits have discovered few novel associations – partly due to the higher incidence of rare alleles – there some exceptions, e.g. for age-related[51] and eye diseases[50] and for blood corpuscle measurements[52].

Second, the excess of partial and overdominant QTLs compared to complete dominant QTLs suggests that heterosis may be caused by polygenic heterozygote advantage (the over-dominance hypothesis) rather than being driven by the superiority of a few dominant alleles over deleterious recessive alleles (the dominance hypothesis). Heterosis – which is closely related to dominance [89, 90] – is of great importance in animal improvement and was the motivation behind breeding the F2 pig population used here.

Third, integrated modelling of dominance across physiological traits and gene and transcript expression traits can suggests causal mechanisms. In some cases we can infer the relationship between a trait and genetic variant is mediated via the expression of a particular transcript rather than by overall gene expression (Fig. 7, Supplementary Fig. S7-1 and Supplementary Table S13). Supplementary Figures S8-1-3 show examples of potential causal links between gene expression and physiological traits, based on co-localisation of dominance eQTLs and QTLs, in each species. Whilst we cannot not prove causality in these examples, selecting on dominance allows us to exclude, for example, additive eQTLs underlying dominance physiological QTLs, which are less likely to be causal. We conjecture that in some cases physiological dominance arises from the complex genetic control of alternative transcripts. Genetic control of alternative splicing has been reported in humans but without reference to dominance[91].

We observed a clear linkage between cis vs trans eQTLs and additive vs dominant eQTLs. Specifically, most dominant eQTLs are trans-acting, and most cis-eQTLs are additive. A potential explanation is that cis-acting causal variants tend to lie in the binding regions of regulatory elements of target genes, and the degree of binding is controlled by local sequence variation, thereby causing additive changes in transcriptional levels. In contrast, causal trans-acting variants are likely to be in or near distant transcription factors (TFs) that regulate the target gene. This could lead to non-additive relationships between TF concentrations and gene expression[92]. If the two chromosomes compete for a limited supply of the TF, then non-additive expression may emerge[93]. In contrast, where a trans-eQTL behaves additively, it may control the expression of a TF which binds to both chromosomes with equal efficiency, so that expression of the target gene is proportional to the amount of TF produced e.g. nuclear factor-κB [94]. Antagonistic pleiotropy might also explain over-dominant trans-acting eQTLs, where heterozygotes express more transcriptional outputs compared to either homozygote[95]. Additionally, long-range physical interactions between promotors and enhancers[96], and the silencing effects of some trans-eQTLs[74] could also produce transcriptional nonlinearity.

Taken together, the consideration of dominance uncovers important and interesting biology in quantitative genetics. It requires only a slightly different analysis workflow, such as that implemented in the ADDO package used here. Even though the greater part of trait heritability is usually additive, there are few if any disadvantages to searching for dominance and therefore we recommend its routine use.

## METHODS

### 1. Ethics Statement

All the data used in this study was collected in three earlier studies, each of which was approved by the corresponding local animal ethical committee, as described for F2 pigs[75], HS rats[76] and HS mice[77].

### 2. Data processing of genotypes, physiological and expression traits in F2 pigs

The F2 pigs were established by crossing Chinese Erhualian and White Duroc pigs in 1998, as described [97]. Genotypes of 1,005 F2 pigs were measured using the porcine SNP60 Beadchip (Illumina) at 62,613 SNP sites, which were filtered by minor allele frequency (MAF) < 0.05, missing rate > 0.1 and minimum frequency of the rarest genotype at each locus > 10, to leave 39,298 SNPs for downstream analysis. A total of 253 complex traits were measured for growth, fatness, meat quality, basal hematology and serum biochemistry. For each trait, we controlled for outliers by removing values more than 5 s.d. from the mean. Gene expression from 493 liver and 583 muscle samples of the same pigs was measured using digital gene expression (DGE), processed into transcript per million (TPM) values for 15,684 and 17,822 transcripts in liver and muscle respectively. The TPM values were quantile normalized and also adjusted for sex, batch and the first ten principal components of expression data.

### 3. Data processing of genotypes, physical and expressional traits in HS rats

The HS rats were descended from eight inbred strains (ACI/N, BN/SsN, BUF/N, F344/N, M520/N, MR/N, WKY/N and WN/N) [98], by rotational breeding over many generations, such that each HS rat chromosome is a mosaic of the founder genomes. In total, the genotypes of 1,407 individuals [76] were measured by a custom Affymetrix array for 257,868 SNPs, as well as a comprehensive measurement of 220 complex traits, including various complex traits related to psychology, basal hematology, basal immunology and serum biochemistry. Transcriptome RNA sequencing (RNA-seq) for 205 amygdala and 192 heart previously unpublished samples from the same animals were analysed in this study. We used different pipelines to quantify gene and transcript expression levels, using the same reference (*Rattus norvegicus* genome, Ensembl RGSC3.4) for consistency with the version used for the array genotypes. For the gene expression levels, we first aligned clean reads to the rat RGSC3.4 reference genome using STAR (v.2.5.3a), and removed duplicated reads by Picard (v.2.5.0). Next, we estimated the raw read counts of each gene using featureCounts (v.1.5.2), and normalized the counts using the Trimmed Mean of M-value (TMM) method, implemented in edgeR (v.3.20.9). For the transcript levels, we used Kallisto (v.0.43.1) to estimate the transcript per million (TPM) of all transcripts in rat RGSC3.4 genome, followed by the quantile normal transformation before GWAS analysis.

### 4. Data processing of genotypes, physical and expressional traits in HS mice

The HS mice originated from eight inbred progenitors (A/J, AKR/J, BALBc/J, CBA/J, C3H/HeJ, C57BL/6J, DBA/2J and LP/J) [99] with a similar design as for HS rats. In total, the genotypes of 2,002 individuals were measured by a custom Illumina assay for 10,168 SNPs [77]. In total 125 traits were measured as described in [100], including basal immunological, basal hematology and models of human disease related to anxiety, asthma, diabetes and obesity. Gene expression data (371 hippocampus samples, 227 liver samples and 197 lung samples) were also measured using Illumina microarray-based assays[79].

### 5. Estimation of additive and dominance variance components

We estimated additive and dominance variance components and heritabilities using GCTA[62] in a standard workflow: We first corrected the raw phenotypes regressing out covariates using lm() function in R. We then standardised the residuals by mean and standard error. We generated ***K***_***a***_, ***K***_***d***_ the additive and dominant Genetic Relationship Matrices (GRMs) of all individuals by GCTA. Pairs of individuals with absolute additive genetic correlation > 0.7 were randomly downsampled to single individuals, and the GRMs rebuilt using the remaining individuals. Finally, we calculated the variance components for additive and dominance effects using GCTA.

### 6. Detection and classification of QTLs and eQTLs

All QTL and eQTL mapping was done using the ADDO[18] toolkit. We fitted three mixed models, namely the Add-Dom (AD), Add (A) and Dom (D) models to detect, classify and compare additive and dominance QTLs. Mixed models correct for unequal relatedness between individuals and to avoid false positive QTL calls. In brief, we model the phenotypic variance covariance matrix 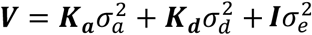 where 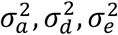 are the additive, dominant and environmental variance components estimated by GCTA. We multiple the phenotype vector and fixed effects design matrix by the matrix ***V*^−0.5^** to convert the mixed model to ordinary least squares with iid errors. To test association as a specific SNP with genotypes AA/AB/BB, we consider the following linear model fixed effect design matrices:

1. “Add (A) Model”: The three genotypes are recoded as 0/1/2 within each locus, ie, as additive genotype dosages, in order to model additive genetic effects. The design matrix is thus a column of ones (for the intercept) and a column of genotype dosages. Except that the variance matrix incorporates dominance effects, the A model is equivalent to the usual additive model used in mixed model GWAS.
2. “Dom (D) Model”: Genotypes AA/AB/BB are coded as heterozygote dosages 0/1/0, to detect loci where the effect of heterozygote AB is from the mean effect of two homozygotes AA and BB. The design matrix is a column of ones and a column of heterozygote dosages.
3. “Add-Dom (AD) Model”: Genotypes are coded as two columns 0/1/2 (additive dosage) and 0/1/0 (heterozygote dosage). The design matrix is a column of ones and both of these columns, and which can model any type of dominance effect by suitable choice of the regression coefficients *β_a_*, *β_d_* corresponding to the additive and dominance columns respectively. We computed the T-statistics *t_Add_*, *t_Dom_* for these coefficients by dividing each by their estimated standard errors.
4. “AvsAD Model”: We used ANOVA to compare the A-Model and AD-Model, to detect loci with significant non-additive effect. Statistical significance was reported as the negative base 10 log p-value of the ANOVA comparison of the models.

We used two p-value thresholds to report significant QTLs: (1) a suggestive threshold (1/N_SNP_), where N_SNP_ is the number of SNPs. Under the null hypothesis where not SNPs are associated one false positive is expected per genome scan; (2) whole genome-wide significance (0.05/N_SNP_), in order to control for false positives caused by multiple tests, and where a false positive should occur once in every 20 GWAS. The width of each QTL was determined using 2 point LOD drop from the peak SNP at the QTL.

We classified each QTL into a dominance type (A: additive/PD: partial dominant/CD: complete dominant/OD: over dominant), based on the log ratio of T-statistics from the Add-Dom Model, i.e. *T* = log_2_ | *t_Dom_*⁄*t_Add_*|, then all the significant QTLs could be classified into four groups using the rules A: (T<log_2_(0.2)), PD: (log_2_(0.2)<T<log_2_(0.8)), CD: (log_2_(0.8)<T<log_2_(1.2)) or OD (T>log_2_(1.2)).

eQTLs were classified as cis-acting if (1) the eQTL localizes in the same chromosome with its target gene (2) the minimum of left and right boundary distances between “Peak SNP” and “gene physical range” < 2Mb, otherwise, it was classified as trans-acting.

## Supporting information

Supplemental Tables zipped

## Acknowledgments

We acknowledge Dr. Amelie Baud, Dr. Na Cai, Dr. Michael Scott, Prof. Dallas Swallow, Prof. Adam S. Wilkins, Prof. David Curtis for their helpful advice and suggestions. The authors also want to thank the editor and reviewers to their practical comments, which contributed to improving the manuscript.

## Data availability

The rat RNAseq data have been deposited at ENA Biostudies under accession numbers E-MTAB-12701, E-MTAB-12693. All other data used in this study are from previously published studies (pig[79, 101], rat[76], mouse[77, 79]) and available as described.

## Code availability

All the codes used for variance decomposition, QTL mapping, RNA-seq, eQTL mapping, variants annotation as well as the respective quality control of raw phenotype and genotype, results summary, comparison and visualization scripts are available at the GitHub website (https://github.com/LeileiCui/Dominance_3Stocks).

## Author contributions

LH, RM and YB conceived and supervised the project. LC performed the analyses and produces the figures. LC and MR drafted the manuscript. LH, MR and YB contributed to the results interpretation and manuscript revision. LH, SX, JG and YB contributed to the pig data used in study. MR, JF, DG, MM, AD, SS, RLA, CM, AFT contributed the rat data used in the study. MR, JF and AB contributed the mouse data used in the study. All authors read and approved the final manuscript.

## References

1. Wright S: Physiological and Evolutionary Theories of Dominance. The American Naturalist 1934, 68:24–53.

2. Kacser H, Burns JA: The molecular basis of dominance. Genetics 1981, 97:639–666.

3. Wilkie AO: The molecular basis of genetic dominance. J Med Genet 1994, 31:89–98.

4. Qian W, Zhang J: Gene dosage and gene duplicability. Genetics 2008, 179:2319–2324.

5. Huber CD, Durvasula A, Hancock AM, Lohmueller KE: Gene expression drives the evolution of dominance. Nature Communications 2018, 9:2750.

6. Sijacic P, Wang W, Liu Z: Recessive antimorphic alleles overcome functionally redundant loci to reveal TSO1 function in Arabidopsis flowers and meristems. PLoS Genet 2011, 7:e1002352.

7. Boettcher S, Miller PG, Sharma R, McConkey M, Leventhal M, Krivtsov AV, Giacomelli AO, Wong W, Kim J, Chao S, et al: A dominant-negative effect drives selection of TP53 missense mutations in myeloid malignancies. Science 2019, 365:599–604.

8. Fertuzinhos S, Legue E, Li D, Liem KF, Jr.: A dominant tubulin mutation causes cerebellar neurodegeneration in a genetic model of tubulinopathy. Sci Adv 2022, 8:eabf7262.

9. Qi H, Zhang H, Zhao Y, Chen C, Long JJ, Chung WK, Guan Y, Shen Y: MVP predicts the pathogenicity of missense variants by deep learning. Nat Commun 2021, 12:510.

10. Carter AJR, Nguyen AQ: Antagonistic pleiotropy as a widespread mechanism for the maintenance of polymorphic disease alleles. BMC Medical Genetics 2011, 12:160.

11. LaFountain AM, Chen W, Sun W, Chen S, Frank HA, Ding B, Yuan YW: Molecular Basis of Overdominance at a Flower Color Locus. G3 (Bethesda) 2017, 7:3947–3954.

12. Merot C, Llaurens V, Normandeau E, Bernatchez L, Wellenreuther M: Balancing selection via life-history trade-offs maintains an inversion polymorphism in a seaweed fly. Nat Commun 2020, 11:670.

13. Fisher RA: The correlation between relatives on the supposition of mendelian inheritance. Trans R Soc Edinb 1918, 53:399–433.

14. Fisher RA: The causes of human variability. The Eugenics review 1919, 10:213–220.

15. Visscher PM, Goddard ME: From R.A. Fisher’s 1918 Paper to GWAS a Century Later. Genetics 2019, 211:1125–1130.

16. Edwards MD, Helentjaris T, Wright S, Stuber CW: Molecular-marker-facilitated investigations of quantitative trait loci in maize : 4. Analysis based on genome saturation with isozyme and restriction fragment length polymorphism markers. Theor Appl Genet 1992, 83:765–774.

17. Li L, Lu K, Chen Z, Mu T, Hu Z, Li X: Dominance, overdominance and epistasis condition the heterosis in two heterotic rice hybrids. Genetics 2008, 180:1725–1742.

18. Cui L, Yang B, Pontikos N, Mott R, Huang L: ADDO: a comprehensive toolkit to detect, classify and visualize additive and non-additive quantitative trait loci. Bioinformatics 2020, 36:1517–1521.

19. Bi W, Kang G, Pounds SB: Statistical selection of biological models for genome-wide association analyses. Methods 2018, 145:67–75.

20. Hall MA, Wallace J, Lucas AM, Bradford Y, Verma SS, Muller-Myhsok B, Passero K, Zhou J, McGuigan J, Jiang B, et al: Novel EDGE encoding method enhances ability to identify genetic interactions. PLoS Genet 2021, 17:e1009534.

21. Li M, Zhang YW, Zhang ZC, Xiang Y, Liu MH, Zhou YH, Zuo JF, Zhang HQ, Chen Y, Zhang YM: A compressed variance component mixed model for detecting QTNs and QTN-by-environment and QTN-by-QTN interactions in genome-wide association studies. Mol Plant 2022, 15:630–650.

22. Gilmour AR, Thompson R, Cullis BR: Average information REML: An efficient algorithm for variance parameter estimation in linear mixed models. Biometrics 1995, 51:1440–1450.

23. Zhang F-T, Zhu Z-H, Tong X-R, Zhu Z-X, Qi T, Zhu J: Mixed Linear Model Approaches of Association Mapping for Complex Traits Based on Omics Variants. Scientific Reports 2015, 5:10298.

24. Wellmann R, Bennewitz J: Bayesian models with dominance effects for genomic evaluation of quantitative traits. Genet Res (Camb) 2012, 94:21–37.

25. Marchini J, Howie B, Myers S, McVean G, Donnelly P: A new multipoint method for genome-wide association studies by imputation of genotypes. Nat Genet 2007, 39:906–913.

26. Gonzalez JR, Armengol L, Sole X, Guino E, Mercader JM, Estivill X, Moreno V: SNPassoc: an R package to perform whole genome association studies. Bioinformatics 2007, 23:644–645.

27. Li Y, Gao Y, Kim YS, Iqbal A, Kim JJ: A whole genome association study to detect additive and dominant single nucleotide polymorphisms for growth and carcass traits in Korean native cattle, Hanwoo. Asian-Australas J Anim Sci 2017, 30:8–19.

28. Hiltpold M, Niu G, Kadri NK, Crysnanto D, Fang ZH, Spengeler M, Schmitz-Hsu F, Fuerst C, Schwarzenbacher H, Seefried FR, et al: Activation of cryptic splicing in bovine WDR19 is associated with reduced semen quality and male fertility. PLoS Genet 2020, 16:e1008804.

29. Doekes HP, Bijma P, Veerkamp RF, de Jong G, Wientjes YCJ, Windig JJ: Inbreeding depression across the genome of Dutch Holstein Friesian dairy cattle. Genet Sel Evol 2020, 52:64.

30. Reynolds EGM, Neeley C, Lopdell TJ, Keehan M, Dittmer K, Harland CS, Couldrey C, Johnson TJJ, Tiplady K, Worth G, et al: Non-additive association analysis using proxy phenotypes identifies novel cattle syndromes. Nat Genet 2021, 53:949–954.

31. Nagai R, Kinukawa M, Watanabe T, Ogino A, Kurogi K, Adachi K, Satoh M, Uemoto Y: Genome-wide detection of non-additive quantitative trait loci for semen production traits in beef and dairy bulls. Animal 2022, 16:100472.

32. Reynolds EGM, Lopdell T, Wang Y, Tiplady KM, Harland CS, Johnson TJJ, Neeley C, Carnie K, Sherlock RG, Couldrey C, et al: Non-additive QTL mapping of lactation traits in 124,000 cattle reveals novel recessive loci. Genet Sel Evol 2022, 54:5.

33. Jiang J, Ma L, Prakapenka D, VanRaden PM, Cole JB, Da Y: A Large-Scale Genome-Wide Association Study in U.S. Holstein Cattle. Front Genet 2019, 10:412.

34. Yang W, Wu J, Yu J, Zheng X, Kang H, Wang Z, Zhang S, Zhou L, Liu J: A genome-wide association study reveals additive and dominance effects on growth and fatness traits in large white pigs. Anim Genet 2021, 52:749–753.

35. Stratz P, Schmid M, Wellmann R, Preuss S, Blaj I, Tetens J, Thaller G, Bennewitz J: Linkage disequilibrium pattern and genome-wide association mapping for meat traits in multiple porcine F2 crosses. Anim Genet 2018, 49:403–412.

36. Lopes MS, Bastiaansen JW, Harlizius B, Knol EF, Bovenhuis H: A genome-wide association study reveals dominance effects on number of teats in pigs. PLoS One 2014, 9:e105867.

37. Coster A, Madsen O, Heuven HC, Dibbits B, Groenen MA, van Arendonk JA, Bovenhuis H: The imprinted gene DIO3 is a candidate gene for litter size in pigs. PLoS One 2012, 7:e31825.

38. Estrada-Reyes ZM, Rae DO, Mateescu RG: Genome-wide scan reveals important additive and non-additive genetic effects associated with resistance to Haemonchus contortus in Florida Native sheep. Int J Parasitol 2021, 51:535–543.

39. Tarsani E, Kranis A, Maniatis G, Avendano S, Hager-Theodorides AL, Kominakis A: Deciphering the mode of action and position of genetic variants impacting on egg number in broiler breeders. BMC Genomics 2020, 21:512.

40. Marchesi JAP, Ono RK, Cantao ME, Ibelli AMG, Peixoto JO, Moreira GCM, Godoy TF, Coutinho LL, Munari DP, Ledur MC: Exploring the genetic architecture of feed efficiency traits in chickens. Sci Rep 2021, 11:4622.

41. M. D. Edwards, C. W. Stuber, Wendel JF: Molecular-Marker-Facilitated Investigations of Quantitative-Trait Loci in Maize. I. Numbers, Genomic Distribution and Types of Gene Action. Genetics 1987, 116:113–125.

42. Vikal Y, Kaur A, Jindal J, Kaur K, Pathak D, Garg T, Singh A, Singh P, Yadav I: Identification of genomic regions associated with shoot fly resistance in maize and their syntenic relationships in the sorghum genome. PLoS One 2020, 15:e0234335.

43. Galli G, Alves FC, Morosini JS, Fritsche-Neto R: On the usefulness of parental lines GWAS for predicting low heritability traits in tropical maize hybrids. PLoS One 2020, 15:e0228724.

44. Monir MM, Zhu J: Dominance and Epistasis Interactions Revealed as Important Variants for Leaf Traits of Maize NAM Population. Front Plant Sci 2018, 9:627.

45. Beukert U, Liu G, Thorwarth P, Boeven PHG, Longin CFH, Zhao Y, Ganal M, Serfling A, Ordon F, Reif JC: The potential of hybrid breeding to enhance leaf rust and stripe rust resistance in wheat. Theor Appl Genet 2020, 133:2171–2181.

46. Bonnafous F, Fievet G, Blanchet N, Boniface MC, Carrere S, Gouzy J, Legrand L, Marage G, Bret-Mestries E, Munos S, et al: Comparison of GWAS models to identify non-additive genetic control of flowering time in sunflower hybrids. Theor Appl Genet 2018, 131:319–332.

47. Seymour DK, Chae E, Grimm DG, Martin Pizarro C, Habring-Muller A, Vasseur F, Rakitsch B, Borgwardt KM, Koenig D, Weigel D: Genetic architecture of nonadditive inheritance in Arabidopsis thaliana hybrids. Proc Natl Acad Sci U S A 2016, 113:E7317–E7326.

48. Feng C, Yi H, Yang L, Kang M: The genetic basis of hybrid male sterility in sympatric Primulina species. BMC Evolutionary Biology 2020, 20:49.

49. Powell JE, Henders AK, McRae AF, Kim J, Hemani G, Martin NG, Dermitzakis ET, Gibson G, Montgomery GW, Visscher PM: Congruence of additive and non-additive effects on gene expression estimated from pedigree and SNP data. PLoS Genet 2013, 9:e1003502.

50. Pozarickij A, Williams C, Guggenheim JA, and the UKBE, Vision C: Non-additive (dominance) effects of genetic variants associated with refractive error and myopia. Mol Genet Genomics 2020, 295:843–853.

51. Guindo-Martínez M, Amela R, Bonàs-Guarch S, Puiggròs M, Salvoro C, Miguel-Escalada I, Carey CE, Cole JB, Rüeger S, Atkinson E, et al: The impact of non-additive genetic associations on age-related complex diseases. Nature Communications 2021, 12:2436.

52. Palmer DS, Zhou W, Abbott L, Baya N, Churchhouse C, Seed C, Poterba T, King D, Kanai M, Bloemendal A, Neale BM: Analysis of genetic dominance in the UK Biobank. bioRxiv 2022:2021.2008.2015.456387.

53. Okbay A, Wu Y, Wang N, Jayashankar H, Bennett M, Nehzati SM, Sidorenko J, Kweon H, Goldman G, Gjorgjieva T, et al: Polygenic prediction of educational attainment within and between families from genome-wide association analyses in 3 million individuals. Nat Genet 2022.

54. Raidan FSS, Porto-Neto LR, Li Y, Lehnert SA, Vitezica ZG, Reverter A: Evaluation of nonadditive effects in yearling weight of tropical beef cattle. J Anim Sci 2018, 96:4028–4034.

55. Akanno EC, Abo-Ismail MK, Chen L, Crowley JJ, Wang Z, Li C, Basarab JA, MacNeil MD, Plastow GS: Modeling heterotic effects in beef cattle using genome-wide SNP-marker genotypes. Journal of Animal Science 2018, 96:830–845.

56. Ertl J, Legarra A, Vitezica ZG, Varona L, Edel C, Emmerling R, Gotz KU: Genomic analysis of dominance effects on milk production and conformation traits in Fleckvieh cattle. Genet Sel Evol 2014, 46:40.

57. Alves K, Brito LF, Baes CF, Sargolzaei M, Robinson JAB, Schenkel FS: Estimation of additive and non-additive genetic effects for fertility and reproduction traits in North American Holstein cattle using genomic information. J Anim Breed Genet 2020, 137:316–330.

58. Serenius T, Stalder KJ, Puonti M: Impact of dominance effects on sow longevity. J Anim Breed Genet 2006, 123:355–361.

59. Su G, Christensen OF, Ostersen T, Henryon M, Lund MS: Estimating additive and non-additive genetic variances and predicting genetic merits using genome-wide dense single nucleotide polymorphism markers. PLoS One 2012, 7:e45293.

60. Lopes MS, Bastiaansen JW, Janss L, Knol EF, Bovenhuis H: Estimation of Additive, Dominance, and Imprinting Genetic Variance Using Genomic Data. G3 (Bethesda) 2015, 5:2629–2637.

61. Costa EV, Diniz DB, Veroneze R, Resende MD, Azevedo CF, Guimaraes SE, Silva FF, Lopes PS: Estimating additive and dominance variances for complex traits in pigs combining genomic and pedigree information. Genet Mol Res 2015, 14:6303–6311.

62. Zhu Z, Bakshi A, Vinkhuyzen AA, Hemani G, Lee SH, Nolte IM, van Vliet-Ostaptchouk JV, Snieder H, LifeLines Cohort S, Esko T, et al: Dominance genetic variation contributes little to the missing heritability for human complex traits. Am J Hum Genet 2015, 96:377–385.

63. Hivert V, Sidorenko J, Rohart F, Goddard ME, Yang J, Wray NR, Yengo L, Visscher PM: Estimation of non-additive genetic variance in human complex traits from a large sample of unrelated individuals. Am J Hum Genet 2021, 108:786–798.

64. Pazokitoroudi A, Chiu AM, Burch KS, Pasaniuc B, Sankararaman S: Quantifying the contribution of dominance deviation effects to complex trait variation in biobank-scale data. Am J Hum Genet 2021, 108:799–808.

65. Hill WG, Goddard ME, Visscher PM: Data and theory point to mainly additive genetic variance for complex traits. PLoS Genet 2008, 4:e1000008.

66. Guindo-Martinez M, Amela R, Bonas-Guarch S, Puiggros M, Salvoro C, Miguel-Escalada I, Carey CE, Cole JB, Rueger S, Atkinson E, et al: The impact of non-additive genetic associations on age-related complex diseases. Nat Commun 2021, 12:2436.

67. Heyne HO, Karjalainen J, Karczewski KJ, Lemmela SM, Zhou W, FinnGen, Havulinna AS, Kurki M, Rehm HL, Palotie A, Daly MJ: Mono- and biallelic variant effects on disease at biobank scale. Nature 2023, 613:519–525.

68. Gruber JD, Vogel K, Kalay G, Wittkopp PJ: Contrasting properties of gene-specific regulatory, coding, and copy number mutations in Saccharomyces cerevisiae: frequency, effects, and dominance. PLoS genetics 2012, 8:e1002497.

69. Schaefke B, Emerson J, Wang T-Y, Lu M-YJ, Hsieh L-C, Li W-H: Inheritance of gene expression level and selective constraints on trans-and cis-regulatory changes in yeast. Molecular biology and evolution 2013, 30:2121–2133.

70. Zhang X, Cal AJ, Borevitz JO: Genetic architecture of regulatory variation in Arabidopsis thaliana. Genome research 2011, 21:725–733.

71. Meiklejohn CD, Coolon JD, Hartl DL, Wittkopp PJ: The roles of cis-and trans-regulation in the evolution of regulatory incompatibilities and sexually dimorphic gene expression. Genome Research 2014, 24:84–95.

72. Pritchard VL, Viitaniemi HM, McCairns RS, Merilä J, Nikinmaa M, Primmer CR, Leder EH: Regulatory architecture of gene expression variation in the threespine stickleback Gasterosteus aculeatus. G3: Genes, Genomes, Genetics 2017, 7:165–178.

73. Wong ES, Schmitt BM, Kazachenka A, Thybert D, Redmond A, Connor F, Rayner TF, Feig C, Ferguson-Smith AC, Marioni JC: Interplay of cis and trans mechanisms driving transcription factor binding and gene expression evolution. Nature communications 2017, 8:1–13.

74. Kaisaki PJ, Otto GW, Argoud K, Collins SC, Wallis RH, Wilder SP, Yau ACY, Hue C, Calderari S, Bihoreau MT, et al: Transcriptome Profiling in Rat Inbred Strains and Experimental Cross Reveals Discrepant Genetic Architecture of Genome-Wide Gene Expression. G3 (Bethesda) 2016, 6:3671–3683.

75. Cui L, Zhang J, Ma J, Guo Y, Li L, Xiao S, Ren J, Yang B, Huang L: Sexually dimorphic genetic architecture of complex traits in a large-scale F2 cross in pigs. Genet Sel Evol 2014, 46:76.

76. Rat Genome S, Mapping C, Baud A, Hermsen R, Guryev V, Stridh P, Graham D, McBride MW, Foroud T, Calderari S, et al: Combined sequence-based and genetic mapping analysis of complex traits in outbred rats. Nat Genet 2013, 45:767–775.

77. Valdar W, Solberg LC, Gauguier D, Burnett S, Klenerman P, Cookson WO, Taylor MS, Rawlins JN, Mott R, Flint J: Genome-wide genetic association of complex traits in heterogeneous stock mice. Nat Genet 2006, 38:879–887.

78. Zhang J, Cui L, Ma J, Chen C, Yang B, Huang L: Transcriptome analyses reveal genes and pathways associated with fatty acid composition traits in pigs. Anim Genet 2017, 48:645–652.

79. Huang GJ, Shifman S, Valdar W, Johannesson M, Yalcin B, Taylor MS, Taylor JM, Mott R, Flint J: High resolution mapping of expression QTLs in heterogeneous stock mice in multiple tissues. Genome Res 2009, 19:1133–1140.

80. McLaren W, Pritchard B, Rios D, Chen Y, Flicek P, Cunningham F: Deriving the consequences of genomic variants with the Ensembl API and SNP Effect Predictor. Bioinformatics 2010, 26:2069–2070.

81. Matsui T, Mullis MN, Roy KR, Hale JJ, Schell R, Levy SF, Ehrenreich IM: The interplay of additivity, dominance, and epistasis on fitness in a diploid yeast cross. Nature Communications 2022, 13:1463.

82. Duenk P, Calus MPL, Wientjes YCJ, Bijma P: Benefits of Dominance over Additive Models for the Estimation of Average Effects in the Presence of Dominance. G3 (Bethesda) 2017, 7:3405–3414.

83. Wientjes YCJ, Bijma P, Calus MPL, Zwaan BJ, Vitezica ZG, van den Heuvel J: The long-term effects of genomic selection: 1. Response to selection, additive genetic variance, and genetic architecture. Genet Sel Evol 2022, 54:19.

84. Liu T, Luo C, Ma J, Wang Y, Shu D, Qu H, Su G: Including dominance effects in the prediction model through locus-specific weights on heterozygous genotypes can greatly improve genomic predictive abilities. Heredity 2022, 128:154–158.

85. Esfandyari H, Bijma P, Henryon M, Christensen OF, Sorensen AC: Genomic prediction of crossbred performance based on purebred Landrace and Yorkshire data using a dominance model. Genet Sel Evol 2016, 48:40.

86. Wilson S, Zheng C, Maliepaard C, Mulder HA, Visser RGF, van der Burgt A, van Eeuwijk F: Understanding the Effectiveness of Genomic Prediction in Tetraploid Potato. Front Plant Sci 2021, 12:672417.

87. Ramstein GP, Larsson SJ, Cook JP, Edwards JW, Ersoz ES, Flint-Garcia S, Gardner CA, Holland JB, Lorenz AJ, McMullen MD, et al: Dominance Effects and Functional Enrichments Improve Prediction of Agronomic Traits in Hybrid Maize. Genetics 2020, 215:215–230.

88. Vuckovic D, Bao EL, Akbari P, Lareau CA, Mousas A, Jiang T, Chen MH, Raffield LM, Tardaguila M, Huffman JE, et al: The Polygenic and Monogenic Basis of Blood Traits and Diseases. Cell 2020, 182:1214–1231 e1211.

89. Xiao Y, Jiang S, Cheng Q, Wang X, Yan J, Zhang R, Qiao F, Ma C, Luo J, Li W, et al: The genetic mechanism of heterosis utilization in maize improvement. Genome Biol 2021, 22:148.

90. Krieger U, Lippman ZB, Zamir D: The flowering gene SINGLE FLOWER TRUSS drives heterosis for yield in tomato. Nat Genet 2010, 42:459–463.

91. Garrido-Martin D, Borsari B, Calvo M, Reverter F, Guigo R: Identification and analysis of splicing quantitative trait loci across multiple tissues in the human genome. Nat Commun 2021, 12:727.

92. Spitz F, Furlong EE: Transcription factors: from enhancer binding to developmental control. Nat Rev Genet 2012, 13:613–626.

93. Porter AH, Johnson NA, Tulchinsky AY: A New Mechanism for Mendelian Dominance in Regulatory Genetic Pathways: Competitive Binding by Transcription Factors. Genetics 2017, 205:101–112.

94. Giorgetti L, Siggers T, Tiana G, Caprara G, Notarbartolo S, Corona T, Pasparakis M, Milani P, Bulyk ML, Natoli G: Noncooperative interactions between transcription factors and clustered DNA binding sites enable graded transcriptional responses to environmental inputs. Mol Cell 2010, 37:418–428.

95. Qian W, Ma D, Xiao C, Wang Z, Zhang J: The genomic landscape and evolutionary resolution of antagonistic pleiotropy in yeast. Cell Rep 2012, 2:1399–1410.

96. Zuin J, Roth G, Zhan Y, Cramard J, Redolfi J, Piskadlo E, Mach P, Kryzhanovska M, Tihanyi G, Kohler H, et al: Nonlinear control of transcription through enhancer-promoter interactions. Nature 2022, 604:571–577.

97. Guo Y, Mao H, Ren J, Yan X, Duan Y, Yang G, Ren D, Zhang Z, Yang B, Ouyang J, et al: A linkage map of the porcine genome from a large-scale White Duroc x Erhualian resource population and evaluation of factors affecting recombination rates. Anim Genet 2009, 40:47–52.

98. Hansen C, Spuhler K: Development of the National Institutes of Health genetically heterogeneous rat stock. Alcohol Clin Exp Res 1984, 8:477–479.

99. Clearn GE, Wilson J, Meredith WM: The use of isogenic and heterogenic mouse stocks in behavioral research. P. 3–22. In g. Lindzey & d. D. Thiessen (ed.), Contrib. To behavior-genetic. 1970.

100. Solberg LC, Valdar W, Gauguier D, Nunez G, Taylor A, Burnett S, Arboledas-Hita C, Hernandez-Pliego P, Davidson S, Burns P, et al: A protocol for high-throughput phenotyping, suitable for quantitative trait analysis in mice. Mamm Genome 2006, 17:129–146.

101. Chen C, Yang B, Zeng Z, Yang H, Liu C, Ren J, Huang L: Genetic dissection of blood lipid traits by integrating genome-wide association study and gene expression profiling in a porcine model. BMC Genomics 2013, 14:848.

