## Supplementary Figures and Tables Legends for "Dominance is common in mammals and is associated with trans-acting gene expression and alternative splicing"

[1]National Key Laboratory for Pig Genetic Improvement and Production Technology, Jiangxi Agricultural University, Nanchang, China. [2]UCL Genetics Institute, University College London, London, UK. [3]School of Life Sciences, Nanchang University, Nanchang, China. [4]Human Aging Research Institute and School of Life Science, Nanchang University, and Jiangxi Key Laboratory of Human Aging, Jiangxi, China. [5]Centre for Genomic Regulation (CRG), The Barcelona Institute of Science and Technology, Barcelona, Spain. [6]BHF Glasgow Cardiovascular Research Centre, University of Glasgow, Glasgow G12 8TA UK. [7]Cardiovascular and Metabolic Disorders Program, Duke-National University of Singapore Medical School, Singapore, [8]Department of Medicine and Life Sciences, Universitat Pompeu Fabra, Barcelona, Spain. [9]Wellcome Trust Centre for Human Genetics, University of Oxford, Oxford UK. [10] Departamento de Psiquiatría y Medicina Legal, Universitat Autonoma de Barcelona, Spain. [11]Genetics and Genomics of Cardiovascular Diseases Research Group, Max Delbrück Center (MDC) for Molecular Medicine in the Helmholtz Association, Berlin, Germany. [12]DZHK (German Center for Cardiovascular Research) partner site Berlin, Berlin, Germany. [13]Charité Universitätsmedizin Berlin, Berlin, Germany. [14]Department of Psychiatry and Behavioral Sciences, Brain Research Institute, University of California, Los Angeles, Los Angeles, CA, USA.

**<u>Address for correspondence</u>**

Lusheng Huang:,

National Key Laboratory for Pig Genetic Improvement and Production Technology, Jiangxi Agricultural University, Nanchang, 330045, P. R. China

Richard Mott:

UCL Genetics Institute, University College London, London, WC1E 6BT, UK

**Supplementary Tables**

**Table S1**: (S1-1) Summary of datasets in F2 pigs, HS rats and HS mice, together with phenotype and genotype quality control methods applied in QTL/eQTL mapping (S1-2) Trait classes and totals.

**Table S2**: Phenotypic variances with standard errors, due to additive and dominance effects (calculated by genome-wide SNP heritability or from accumulated significant QTLs) of 242 physical traits in F2 pigs (S2-1), 206 physical traits of HS rats (S2-2) and 124 physical traits of HS mice (S2-3), together with average values across each trait class.

**Table S3**: Counts and proportions of significant QTLs under dominance types within each population, and the corresponding missing rate compared to the additive GWAS, under suggestive thresholds and whole genome thresholds.

**Table S4**: Descriptive statistics, effect classification and consequence annotation of all significant QTL detected by AD Model in F2 pigs (S4-1), HS rats (S4-2) and HS mice (S4-3). (Rows with -log$_{10}$(*P*) values of ADvsA Model > 5 are in red, rows where -log$_{10}$(*P*) values of AD Model > 6 are in blue.)

**Table S5**: Summary of novel pleiotropic QTLs detected by AD model within each population. (Different shared QTL regions are separated by grey background and the rows with red numbers indicate novel QTLs where -log$_{10}$(*P*) values of ADvsA Model > 5)

**Table S6**: Summary of homologous QTLs of common physical traits across three stocks. (Different trait groups are separated by frame and QTL rows of different stocks are colored differently)

**Table S7**: Gene expression heritabilities and standard errors explained by additive and dominance effects estimated by genome-wide SNPs within each tissue of F2 pigs (S7-1), HS rats (S7-2) and HS mice (S7-3).

**Table S8**: Counts and proportions of significant eQTLs classified by dominance effect type within each tissue and each population under different significant thresholds (S8-1); Proportions of novel eQTLs detected by AD model compared to A model in cis/trans groups (S8-2).

**Table S9**: Classification of eQTLs detected by AD Model (S9-1, F2 pigs liver tissue; S9-2, F2 pigs muscle tissue; S9-3, HS rats amygdala genes expression; S9-4, HS rats heart genes expression; S9-5, amygdala transcripts expression; S9-6, heart transcripts expression; S9-7, HS mice hippocampus gene expression; S9-8, HS mice liver gene expression; S9-9, HS mice lung gene expression). Rows with -$\log_{10}(P)$ values of ADvsA Model > 6 are in red, remaining rows with -$\log_{10}(P)$ values of AD Model > 10 are in blue.

**Table S10**: Summary of pleiotropic eQTLs (S10-1, F2 pigs liver genes expression; S10-2, F2 pigs muscle genes expression; S10-3, HS rats amygdala genes expression; S10-4, HS rats heart genes expression; S10-5, amygdala transcripts expression; S10-6, heart transcripts expression; S10-7, HS mice hippocampus gene expression; S10-8, HS mice liver gene expression; S10-9, HS mice lung gene expression). Lines in bold indicate cis-eQTLs.

**Table S11**: Summary of eQTLs shared across tissues within each population (S11-1, liver-muscle in F2 pigs; S11-2, amygdala-heart gene expression levels in HS rats; S11-3, amygdala-heart transcript expression levels in HS rats; S11-4, hippocampus-liver in HS mice; S11-5, hippocampus-lung in HS mice; S11-6, liver-lung in HS mice). All rows are ordered by (1) whether from the same gene (2) the -$\log_{10}(P)$ values of ADvsA Model.

**Table S12**: Counts of significant cis-eQTLs and trans-eQTLs under four different effect types within each tissue of three populations under different significant thresholds, together with the correlation test between different genetic modes and different regulatory types (Table S12-1, A/PD/CD/OD vs cis/trans; Table S12-2, A & PD/CD & OD vs cis/trans, using Add-eQTL represents A- & PD-eQTL and Dom-eQTL represents CD- & OD-eQTL).

**Table S13**: Counts of significant eQTLs classified by transcript compositions (one transcript or multi transcripts) and by dominance type, in HS rats heart and amygdala, together with the correlation test between dominance type (A/PD/CD/OD) and transcript compositions

**Table S14**: Summary of shared QTL-eQTL pairs (S14-1, F2 pigs liver tissue; S14-2, F2 pigs muscle tissue; S14-3, HS rats amygdala genes; S14-4, HS rats heart genes; S14-5, amygdala transcripts; S14-6, heart transcripts; S14-7, HS mice hippocampus gene; S14-8, HS mice liver gene; S14-9, HS mice lung gene). Lines in bold indicate the QTL shared with cis-eQTL.

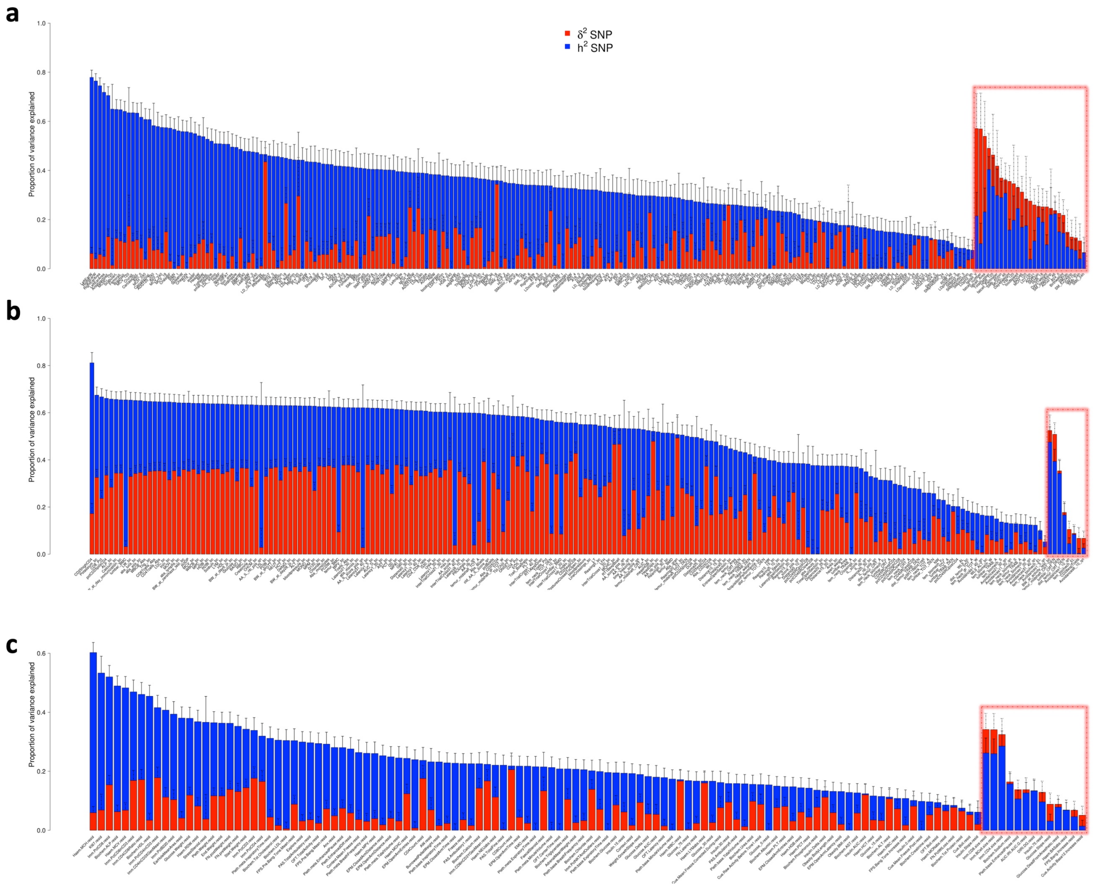


**Fig. S1 Additive (blue bars, $h^2_{SNP}$) and Dominance (red bars, $\delta^2_{SNP}$) Variance Components of all traits across three populations.** (a): F2 pigs, (b): HS rats, (c): HS mice. Each vertical bar represents one trait, labelled on x-axis. Height of each bar represents the total heritability of the corresponding trait, partitioned into additive (blue) and dominance (red) components, as estimated by GCTA. The solid and dashed error bars denote the standard errors of additive and dominance effects in each trait, respectively. Red dashed boxes highlight traits with higher $\delta^2_{SNP}$ than $h^2_{SNP}$, for which are sorted by $\delta^2_{SNP}$ in the red boxes, and other traits are sorted by $h^2_{SNP}$ from left to right.

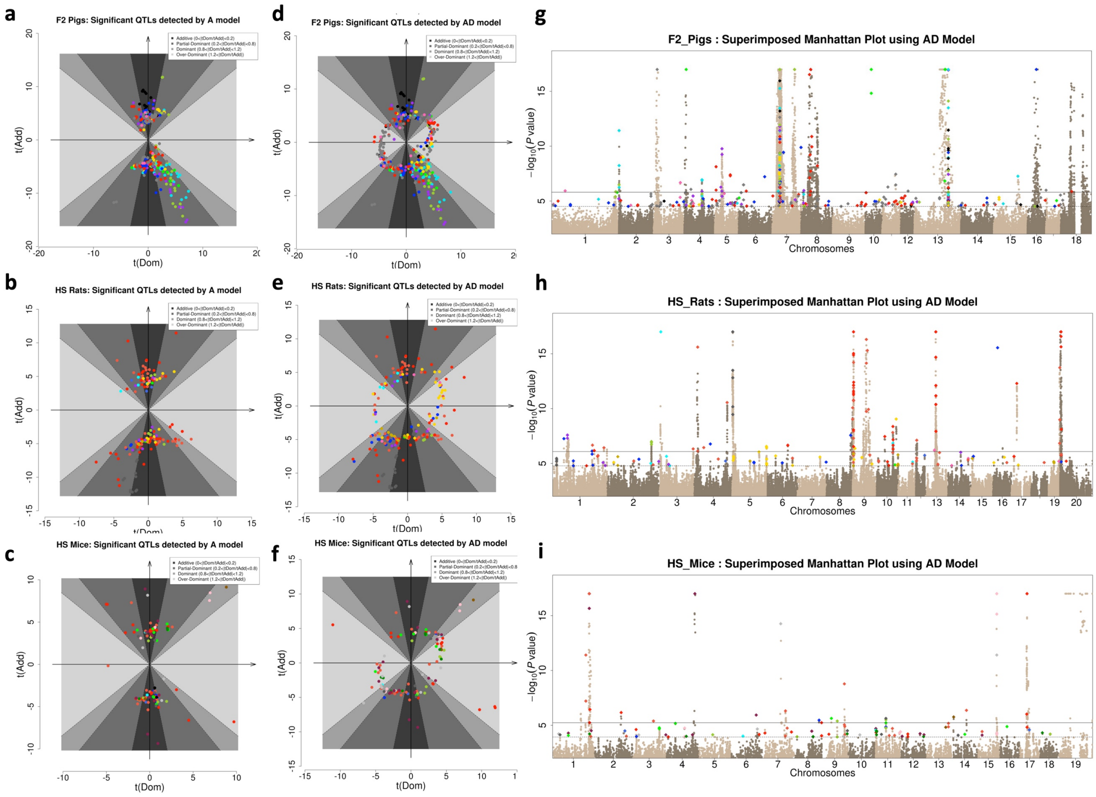


**Fig. S2-1 QTLs detected by AD model.** (a-f) show the same data as in Fig 2 where QTLs are classified according to the ratios of their T-statistics from the AD-model, except that in this figure the ratios have not been log-transformed. Each row is one population (a,b: F2 pigs; c,d: HS rats; e,f: HS mice). (a-c): QTLs detected by A-model. (d-f): QTLs detected by AD-model. Within each panel, the x-axis and y-axis represent $t_{Dom}$ and $t_{Add}$, the standardized dominance ($\beta_{Dom}$) and additive effects ($\beta_{Add}$), respectively. Each dot is one trait, colour-coded by trait type using the same coding as in Fig 2. The dot localizations represent different inheritance models (from deep grey to light grey are additive, partial-dominant, complete-dominant and over-dominant). (g-i) Aggregate Manhattan plots of the AD GWAS of all physiological traits in pig (g), rat (h) and mouse (i). Associations with -log(*P*) > 18 are truncated and data points with -log(*P*) < 2 are not shown. The same coloring scheme is used for the peak QTL of each trait, the solid and dashed horizontal line represents the average suggestive and whole-genome significant thresholds across all traits.

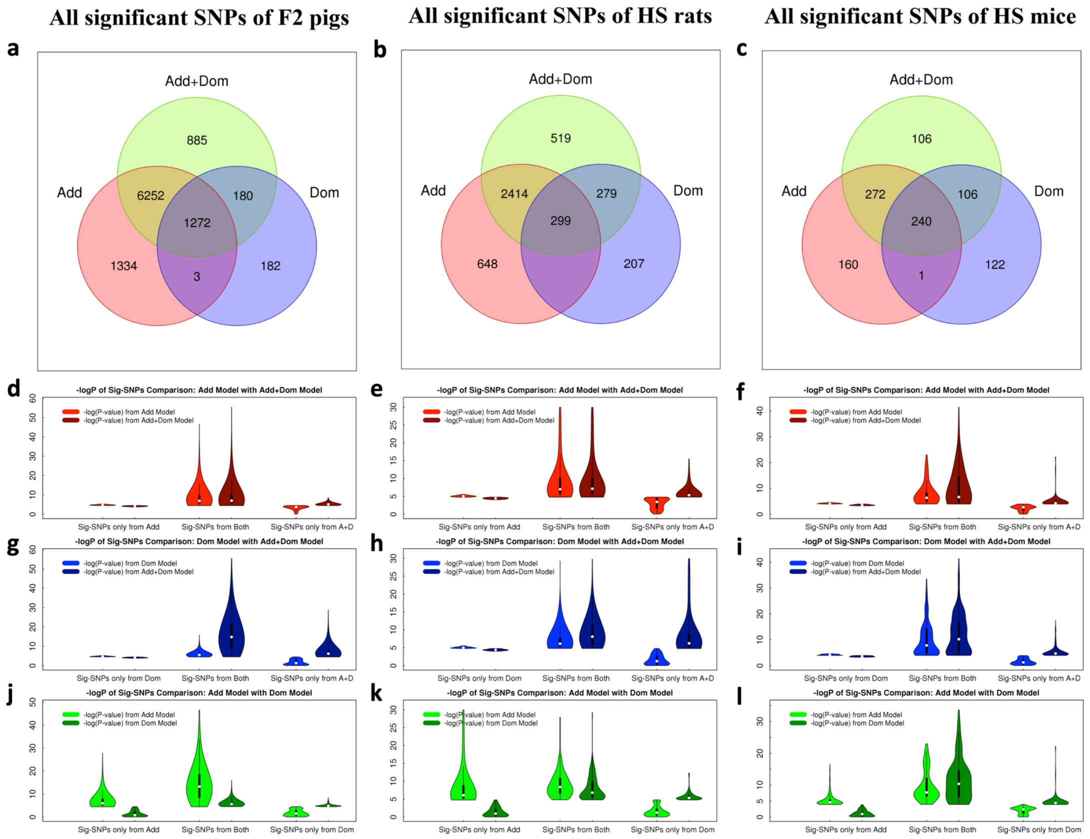


**Fig. S2-2 Comparison of QTL detection among A, D and AD models.** (a-c) Venn diagrams showing overlapping significant SNPs detected by A model (red), D model (blue) and AD model (green) across 247 traits of F2 pigs (a), 207 traits of HS rats (b) and 124 traits of HS mice (c). (d-f) Violin plots showing the distributions of -log(*P*) values of A model and AD model only using SNPs detected by A model (left), SNPs detected by both A and AD models (middle) and SNPs only detected by D model (right) in pig (d), rat (e) and mouse (f), respectively. (g-i) Similar violin plots comparing SNPs detected by D and AD models in pig (g), rat (h) and mouse (i). (j-l) Similar violin plots comparing SNPs detected by A and D models in pig (j), rat (k) and mouse (l).

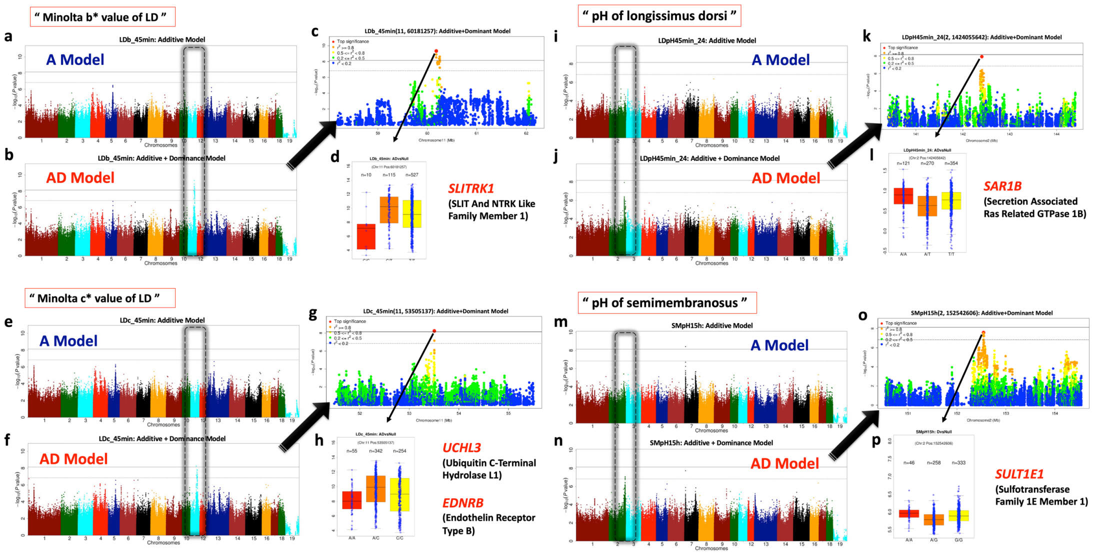


**Fig. S3-1 Over-dominant QTLs in F2 pigs.** (a-d) Manhattan plots of the Minolta b* value of longissimus dorsi muscle based on (a) A model and (b) AD model. (c) regional plot and (d) genotype boxplot showing phenotypic distribution of the peak SNP of AD model. Within each Manhattan plot, the solid and dashed horizontal line show suggestive and whole-genome significant thresholds and the black dashed box indicates the QTL which was only detected by AD model. Also shown are similarly structured plots for other three traits: (e-h) Minolta c* value of longissimus dorsi muscle, (i-l) the pH value of longissimus dorsi muscle, (m-p) pH value of semimembranosus (m-p).

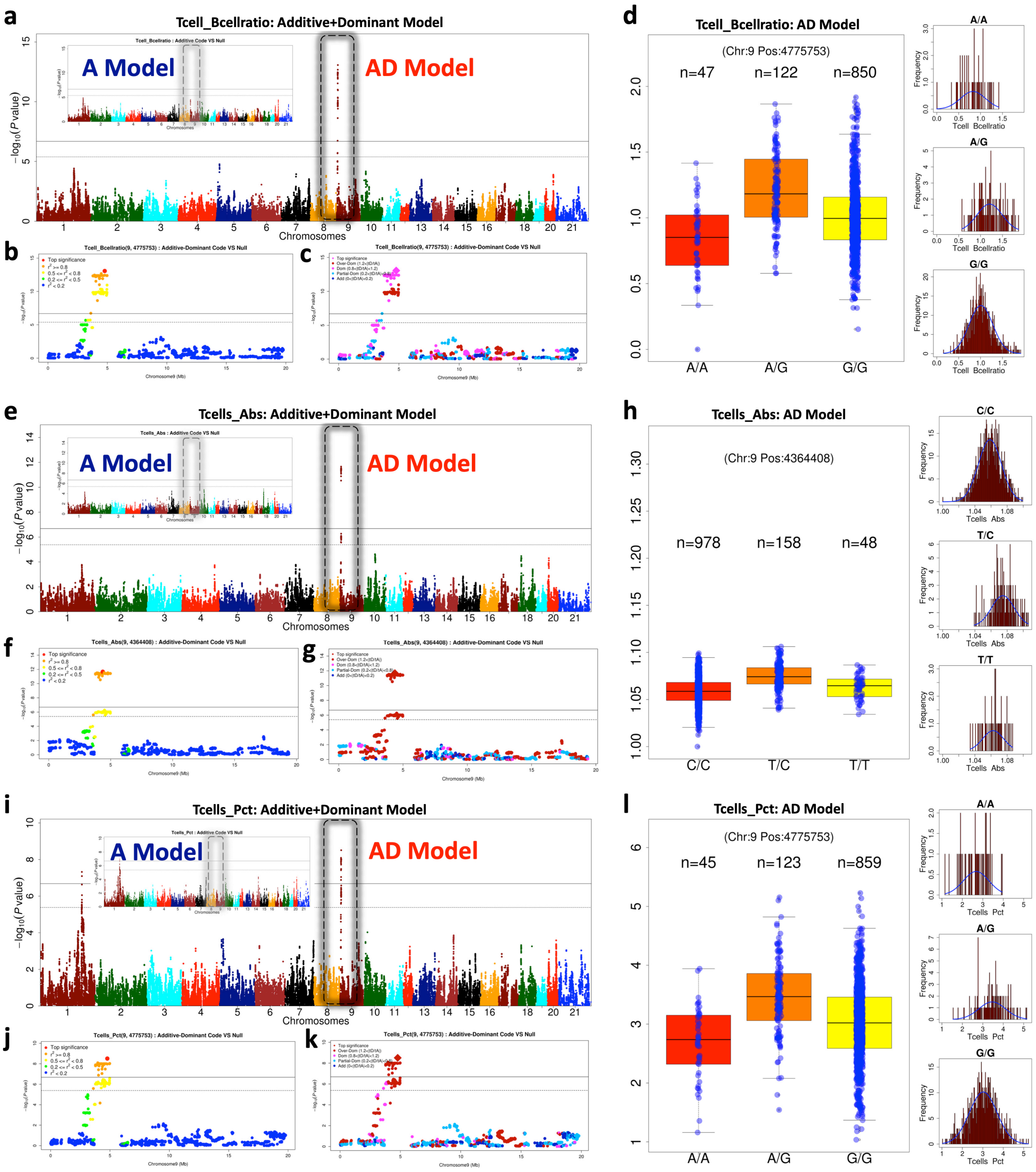


**Fig. S3-2 Over-dominant HS rat QTLs.** (a-d) Manhattan plots of the ratio of T cells to B cells based on A and AD models. Regional Manhattan plots of the peak SNP of AD model showing (b) the LD ($r^2$) with the peak SNP and (c) the different dominance types within the locus. Within each Manhattan plot, the solid and dashed horizontal lines represents suggestive and whole-genome significant thresholds, and the black dashed box highlights the QTL. (d) Boxplot and histogram of phenotypic distributions classified by the peak SNP genotypes. Similar plots shown in (e-h) for absolute T cells counts and (i-l) for proportion of T cells in WBC.

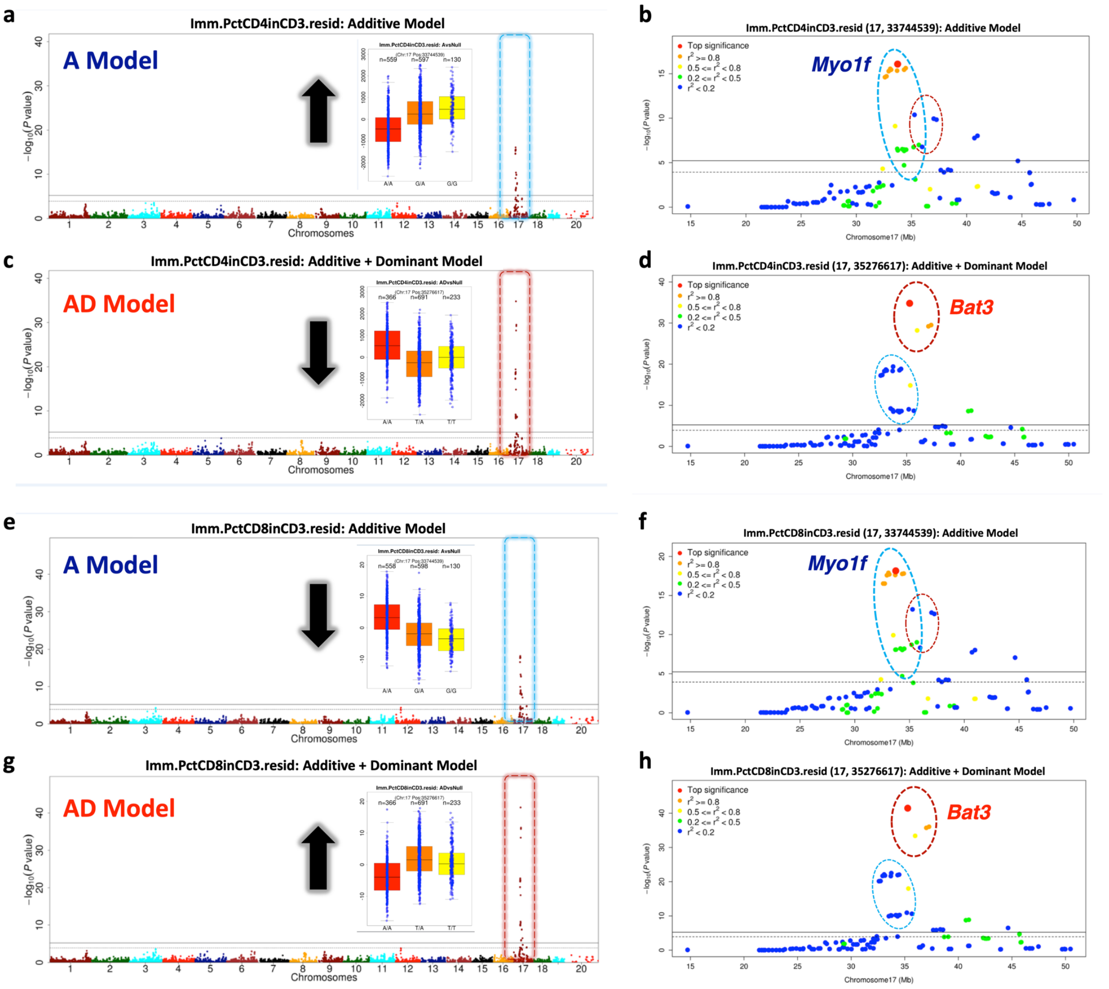


**Fig. S3-3 Over-dominant QTLs implicate a novel causal gene regulating the proportions of CD4+ and CD8+ T cells in HS mice, compared to additive QTL modelling.** (a-d) Manhattan plots and regional Manhattan plots of the proportion of CD4+ T cells in CD3+ T cells based on A model (a, b) and AD model (c, d), respectively. (e-h) Manhattan plots and regional Manhattan plots of the proportion of CD8+ T cells in CD3+ T cells based on A model (e, f) and AD model (g, h). Within each Manhattan plot, the blue and red dashed boxes highlight the QTLs detected by Add model and AD model separately, along with the boxplots of phenotypic distribution of the peak SNPs and black arrows are used to indicate the effect directions of peak SNPs. The blue and red dashed ellipses are used to highlight the linked SNP groups for each model in the regional plots, labelled by the closest annotated genes (*Bat3*: AD Model, *Myo1f*: A Model).

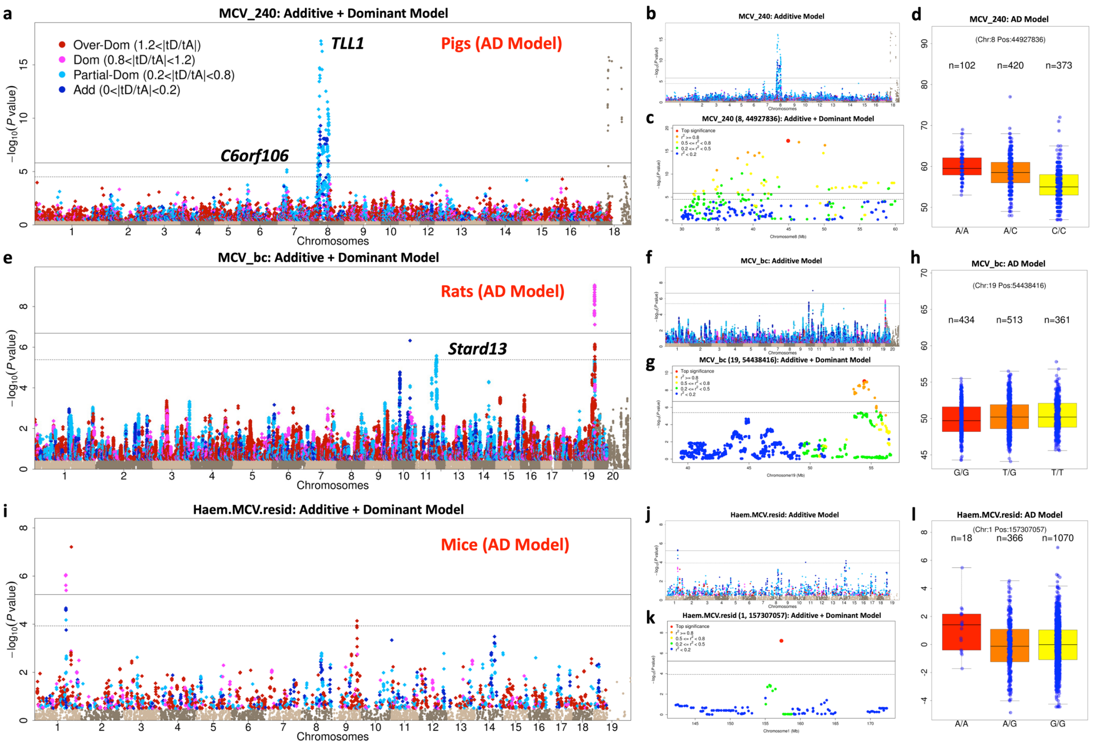


**Fig. S4-1 Homologous QTLs for mean corpuscular volume (MCV).** (a-d) Manhattan plots in F2 pigs based on (a) AD model and (b) A model (c) regional plot (d) phenotypic distribution boxplot of the peak SNP at chr8: 44,927,836 bp of AD model. (e-h) Homologous QTL for HS rats at the peak SNP chr19: 54,438,416 bp. (i-l) Homologous QTL in HS mice at peak SNP chr1: 157,307,057 bp. Within each Manhattan plot, the solid and dashed horizontal line represents the suggestive and whole-genome significance thresholds and all SNPs with -$\log_{10}(P)$ values > 0.5 are coloured according to their dominance classification.

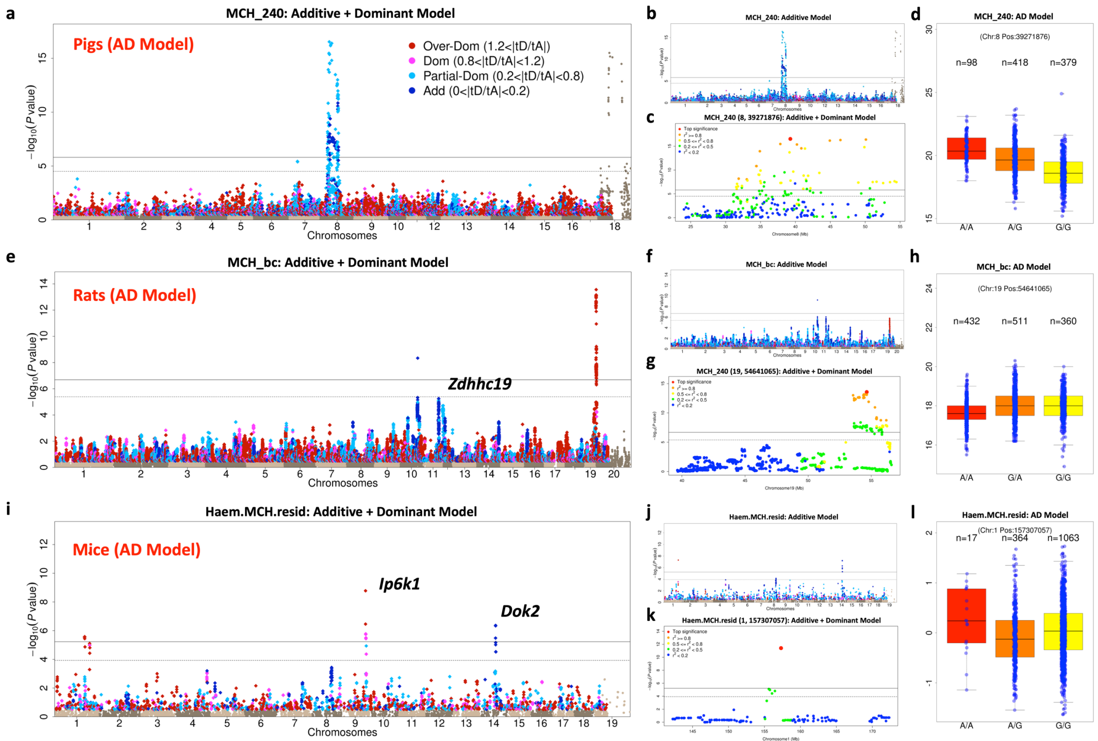


**Fig. S4-2 Homologous QTLs for mean corpuscular volume (MCV).** (a-d) Manhattan plots in F2 pigs based on (a) AD model and (b) A model (c) regional plot (d) phenotypic distribution boxplot of the peak SNP at chr8: 44,927,836 bp of AD model. (e-h) Homologous QTL for HS rats at the peak SNP chr19: 54,438,416 bp. (i-l) Homologous QTL in HS mice at peak SNP chr1: 157,307,057 bp. Within each Manhattan plot, the solid and dashed horizontal line represents the suggestive and whole-genome significance thresholds and all SNPs with -$\log_{10}(P)$ values > 0.5 are coloured according to their dominance classification.

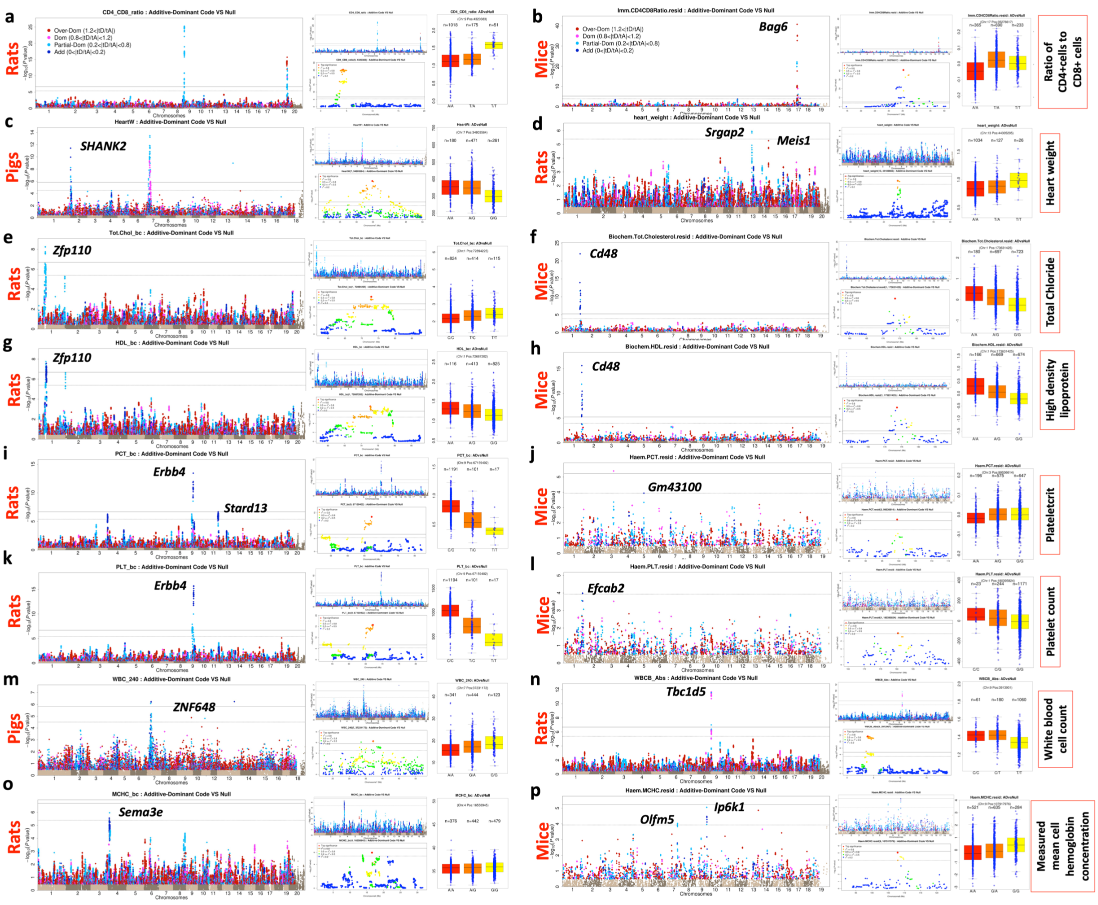


**Fig. S4-3 Homologous QTLs.** (a-b) Ratio of CD4+cells to CD8+ cells in rat and mouse. (c-d) Heart weight of pig and rat. (e-f) Total chloride in rat and mouse. (g-h) High density lipoprotein in rat and mouse. (i-j) Plateletcrit in rat and mouse. (k-l) Platelet count in rat and mouse. (m-n) White blood cell count in pig and rat. (o-p) mean cell hemoglobin concentration on rat and mouse. Within each panel are shown Manhattan plots based on AD model (left) and A model (middle upper) of the specified trait in the population, as well as the regional plot (middle lower) and phenotypic distribution boxplot (right) of the peak SNP in the AD model.

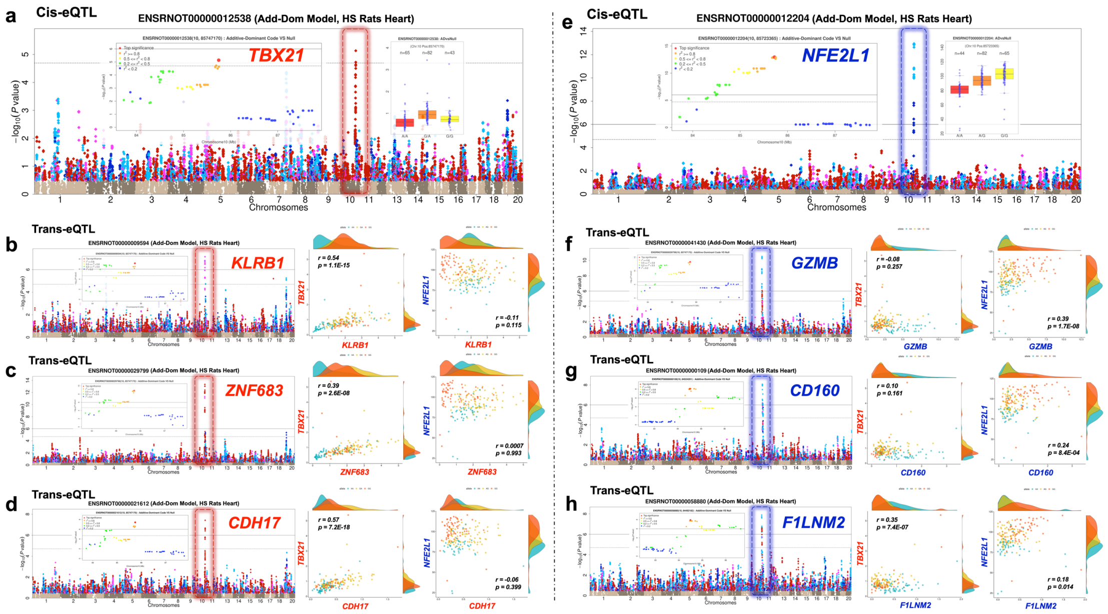


**Fig. S5-1 Dominant cis and trans transcript eQTLs at the hotspot chr10:85Mb-86Mb in HS rat heart.** (a, e) cis-eQTLs with different inheritance modes, (a) *Tbx21* (over-dominant) and (e) *Nf32l1* (partial-dominant). (b-d) Manhattan plots of over-dominant trans-eQTLs for genes *Klrb1*, *Znf683*, *Cdh17*. (f-h) Manhattan plots of partial-dominant trans-eQTLs for genes *Gzmb*, *Cd160*, *F1lnm2*. Within each Manhattan plot, the eQTL is marked by a dotted rectangular frame, with the same colour as the peak SNP dot (Blue - additive; Sky blue - partial-dominant; Purple - complete-dominant; Red - over-dominant), and all linked SNPs with -log10(P) > 0.5 are coloured the same. The regional Manhattan plots of the peak signal of each eQTL and the scatter plots of two cis-eQTLs are also shown as insets. The pairs of scatter plots to the right of each Manhattan plot compare the expression of each trans-eQTL with *Tbx21, Nf32l1,* together with their Pearson correlation coefficients and the p-value of test that the correlation is zero. Each dot represents one animal, colour-coded by the genotype of the peak SNP.

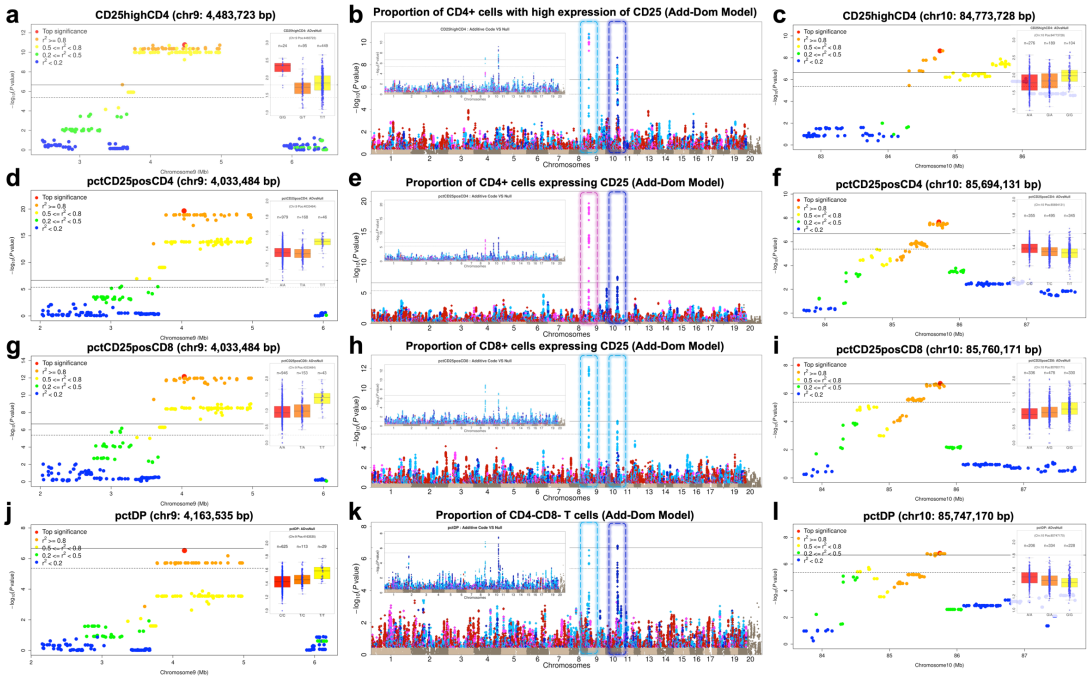


**Fig. S5-2 Interplay of immunology-related QTLs in HS rats.** A locus at chr10: 85Mb-86Mb (cf Fig. 5) is an additive QTL for four rat immunology traits, which are also associated with dominant QTLs at chr9: 4Mb-5Mb**.** Manhattan plots based on AD model for (i) the proportions of CD4+ cells with high expression of CD25 (a-c) (ii) the proportions of CD4+ cells expressing CD25 (d-f) (iii) the proportions of CD8+ cells expressing CD25 (g-i) (iv) the proportions of CD4+CD8- cells (j-l), as well as two corresponding regional Manhattan plots (the left column, the dominant chr9 QTL; the right column, the additive chr10 QTL). Within each Manhattan plot (b, e, h, k), the corresponding GWAS based on A model are shown in the left upper corner.

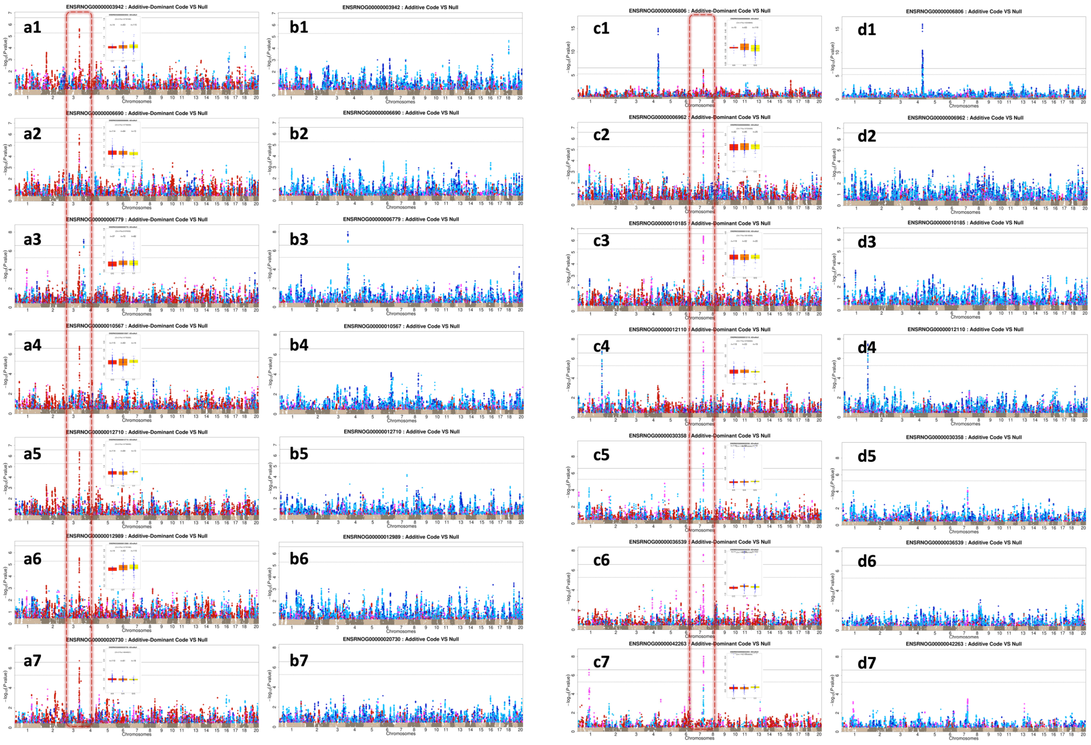


**Fig. S5-3 Dominant eQTLs detected by AD model from two trans-acting hotspots (a1-a7, chr3: 147Mb-149Mb; c1-c7, chr7: 107Mb-108Mb) in HS rat heart.** Left two columns containing the Manhattan plots based on AD model (a1-a7) and A model (b1-b7), (from the top down the genes *Asnsd1*, *Tmem168*, *Crot*, *Angel1*, *Ubac2*, *Serinc2* and *Nbr1*) that are trans-regulated by over-dominant eQTLs. Right two columns show Manhattan plots based on AD model (c1-c7) and A model (d1-d7), (from top down *Setmar*, *Stk32c*, *RGD1306941*, *Col17a1*, *Olr312*, *U6* and *D4A0M7*_RAT). Red dashed boxes indicate the hotspots.

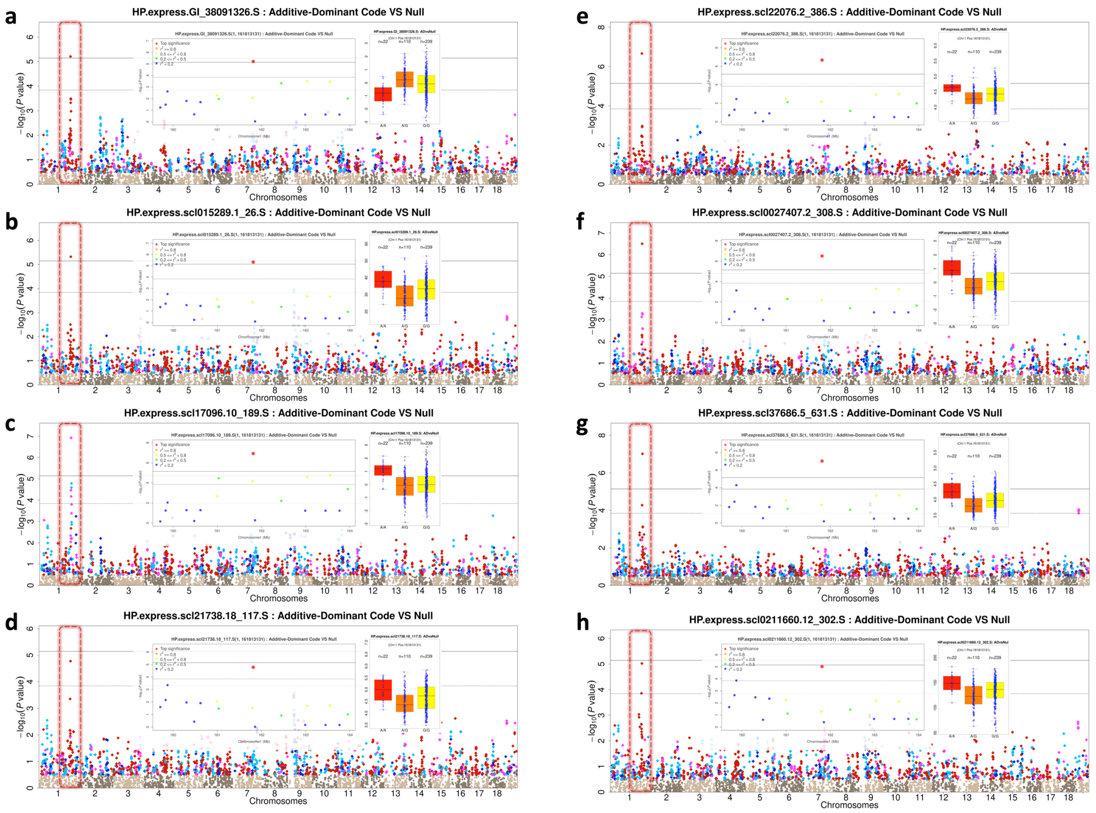


**Fig. S5-4 Dominant eQTLs from a trans-acting hotspot in HS mice hippocampus.** Manhattan plots, regional plots and phenotypic distribution boxplots of the peak SNPs from GWAS across eight genes in HS mice hippocampus. Each of these genes is simultaneously trans-regulated by a dominant cis eQTL located in the hotspot (chr1: 161Mb-163Mb), highlighted by red dashed boxes.

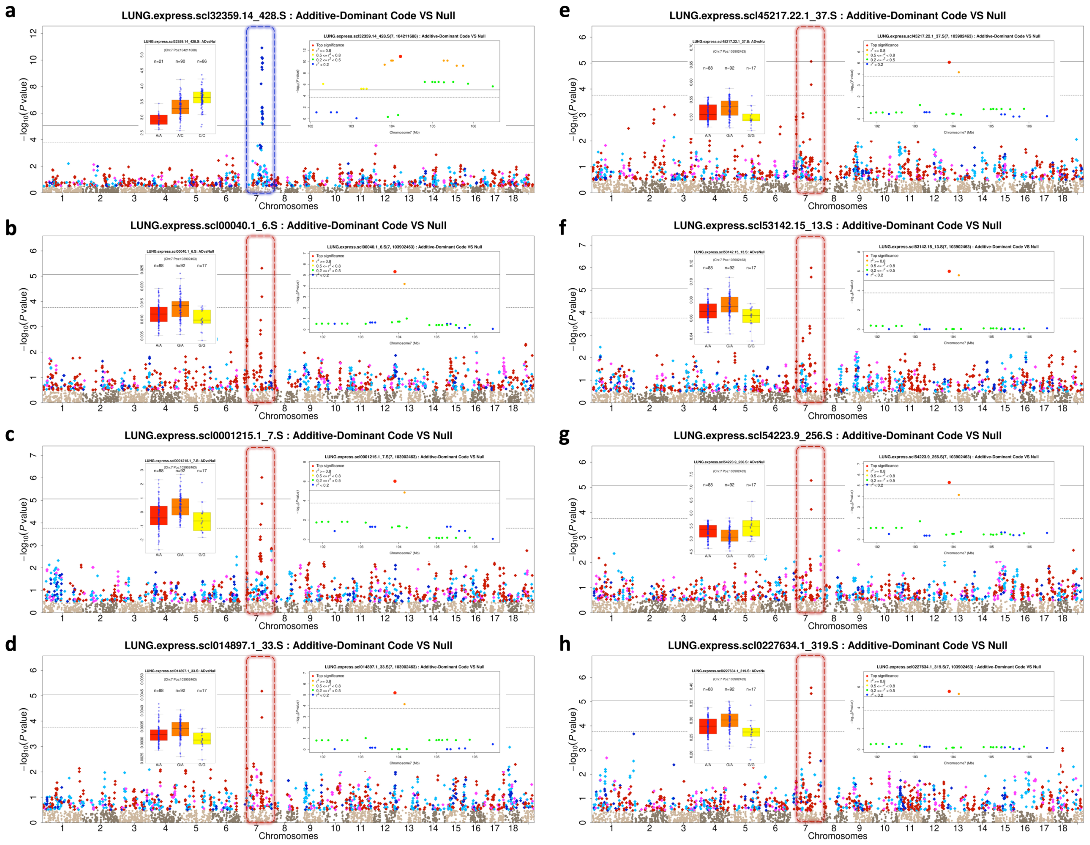


**Fig. S5-5 Dominant eQTLs from a trans-acting hotspot in HS mice lung.** Manhattan plots, regional plots and phenotypic distribution boxplots at the peak SNPs across eight genes in HS mice lung, showing seven genes *Mef2a* (b), *Rassf4* (c), *Trip12* (d), *Dis3* (e), *Entpd1* (f), *Rbmx* (g), *Camsap1* (h) that are simultaneously trans-regulated by an over-dominant cis eQTL for *Nars2* at chr7:103Mb-106Mb (a). Blue and red dashed boxes to highlight the hotspot.

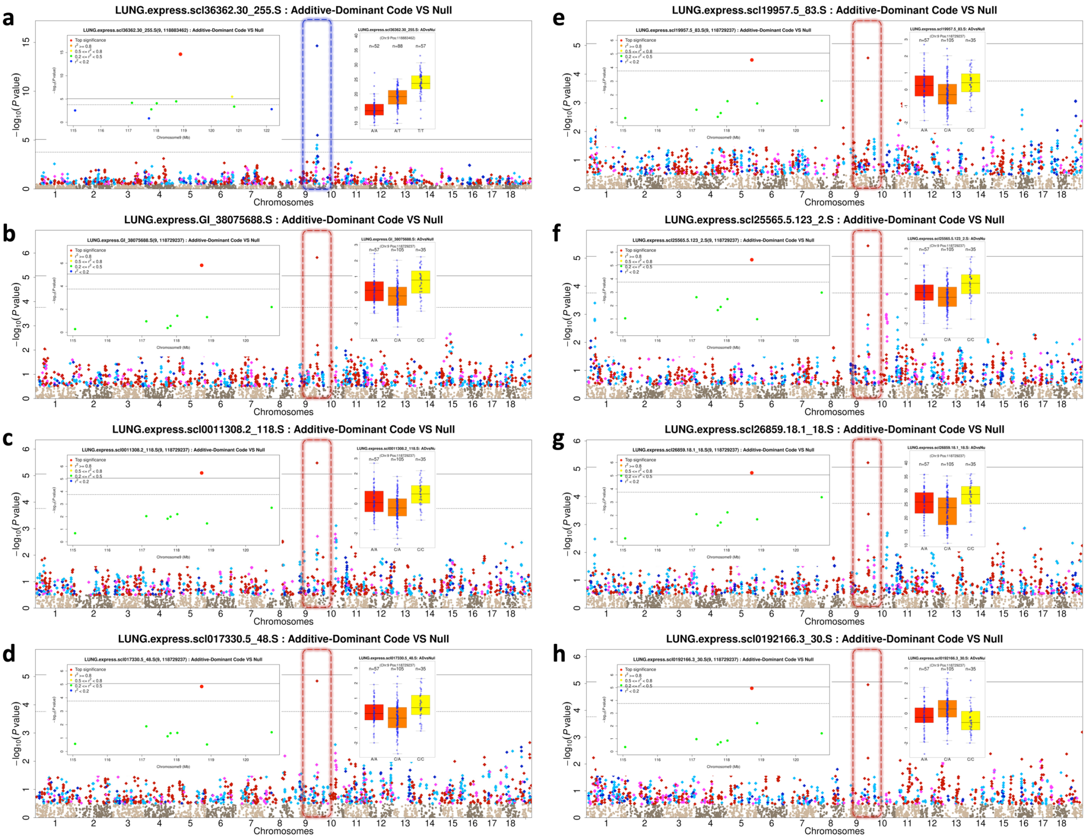


**Fig. S5-6 Dominant eQTLs from a trans-acting hotspot in HS mice lung.** Manhattan plots, regional plots and phenotypic distribution boxplots at the peak SNPs across eight genes in HS mice lung, showing seven genes *AL845491.3* (b), *Abi1* (c), *Minpp1* (d), *Ttpal* (e), *Akirin2* (f), *Ptpn12* (g), *Sardh* (h) that are simultaneously trans-regulated by an over-dominant cis eQTL for *Itga9* at chr9:117Mb-121Mb (a). Blue and red dashed boxes highlight the hotspot.

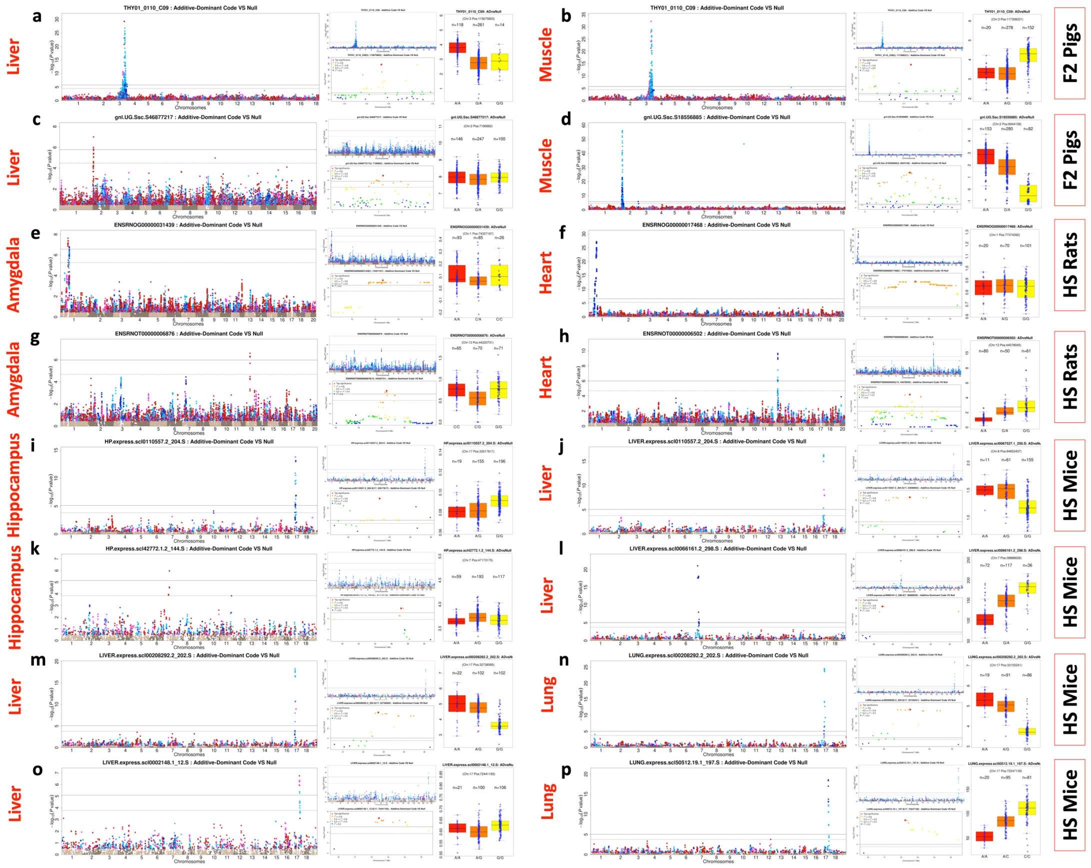


**Fig. S5-7 Tissue-conserved and tissue-specific dominant eQTLs.** Examples of tissue-conserved eQTLs of (1) *GPN1* between liver (a) and muscle (b) in F2 pigs (2) *H2-Q1* between hippocampus (i) and liver (j) in HS mice (3) *9030612M13Rik* between liver (m) and lung (n) in HS mice. Examples of tissue-specific eQTLs shared between (1) *Mgea5* of liver (c) and *Nudt22* of muscle (d) in F2 pigs (2) *ENSRNOG00000031439* of amygdala (e) and *Trappc6a* of heart (f) in HS rats (3) *Nut* of amygdala (g) and *Dyrk3* of heart (h) in HS rats (4) *Dio3* of hippocampus (h) and *Pop4* of liver (l) in HS mice (5) *St8sia5* of liver (o) and *Clip4* of lung (p) in HS mice. Within each panel are shown the Manhattan plots based on AD model (left) and A model (middle upper) as well as the regional plot (middle lower) and phenotypic distribution boxplot (right) of the peak SNP of AD model.

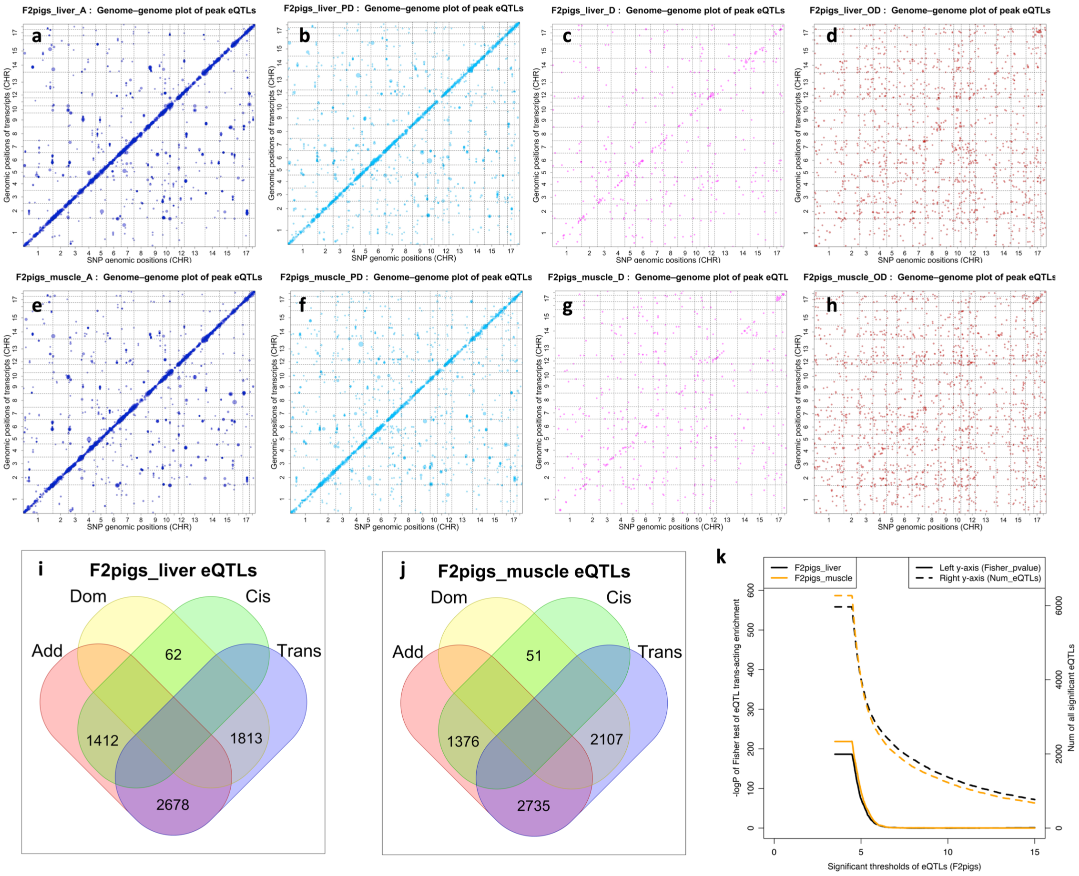


**Fig. S6-1 Trans-acting enrichment among dominant eQTLs in liver and muscle in F2 pigs.** (a-h) eQTL locations of transcript level eQTLs, filtered by dominance type. Each dot represents an eQTL significant at suggestive level (i.e. one false positive expected per transcript). x-axis: eQTL chromosome positions (bp), y-axis: physical gene location. First row (a-d): liver; Second row (e-h): muscle. The four columns represent dominance types (blue: additive A, sky blue: partial-dominant PD, purple: complete-dominant CD, red: over-dominant OD). (i: liver, j: muscle) Venn diagrams comparing eQTL cross-classifications, (1) Generalised Additive eQTLs, (A-eQTLs and PD-eQTLs), (2) Generalised Dominance eQTLs, (CD-eQTLs and OD-eQTLs) vs (3) Cis-acting eQTLs, (4) Trans-acting eQTLs. (k) Dominance trans-acting enrichment (left y-axis, solid lines) and the counts of significant eQTLs (right y-axis, dashed lines) under different -log10(P) eQTL significance thresholds (x-axis). Enrichment is quantified by the -log10(P) values of Fisher exact test between dominance (Add/Dom eQTLs) and regulation types (cis/trans-acting) of significant eQTLs within rat liver (black) and muscle (orange), respectively.

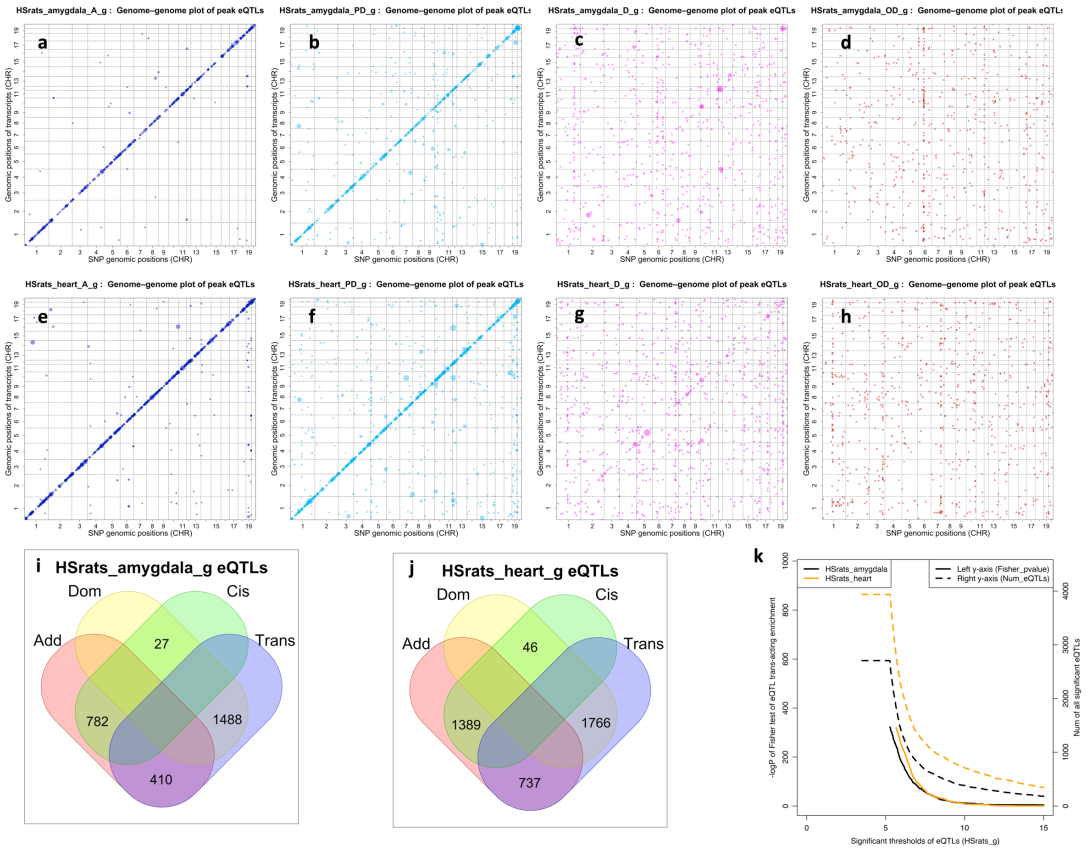


**Fig. S6-2 Trans-acting enrichment among dominant eQTLs of amygdala and heart tissues in HS rats.** (a-h) eQTL locations of gene level eQTLs, filtered by dominance type. Each dot represents an eQTL significant at suggestive level (i.e. one false positive expected per transcript). x-axis: eQTL chromosome positions (bp), y-axis: physical gene location. First row (a-d): amygdala; Second row (e-h): heart. The four columns represent dominance types (blue: additive A, sky blue: partial-dominant PD, purple: complete-dominant CD, red: over-dominant OD). (i: amygdala, j: heart) Venn diagrams comparing eQTL cross-classifications, (1) Generalised Additive eQTLs, (A-eQTLs and PD-eQTLs), (2) Generalised Dominance eQTLs, (CD-eQTLs and OD-eQTLs) vs (3) Cis-acting eQTLs, (4) Trans-acting eQTLs. (k) Dominance trans-acting enrichment (left y-axis, solid lines) and the counts of significant eQTLs (right y-axis, dashed lines) under different -log10(P) eQTL significance thresholds (x-axis). Enrichment is quantified by the -log10(P) values of Fisher exact test between dominance (Add/Dom eQTLs) and regulation types (cis/trans-acting) of significant eQTLs within rat amygdala (black) and heart (orange), respectively.

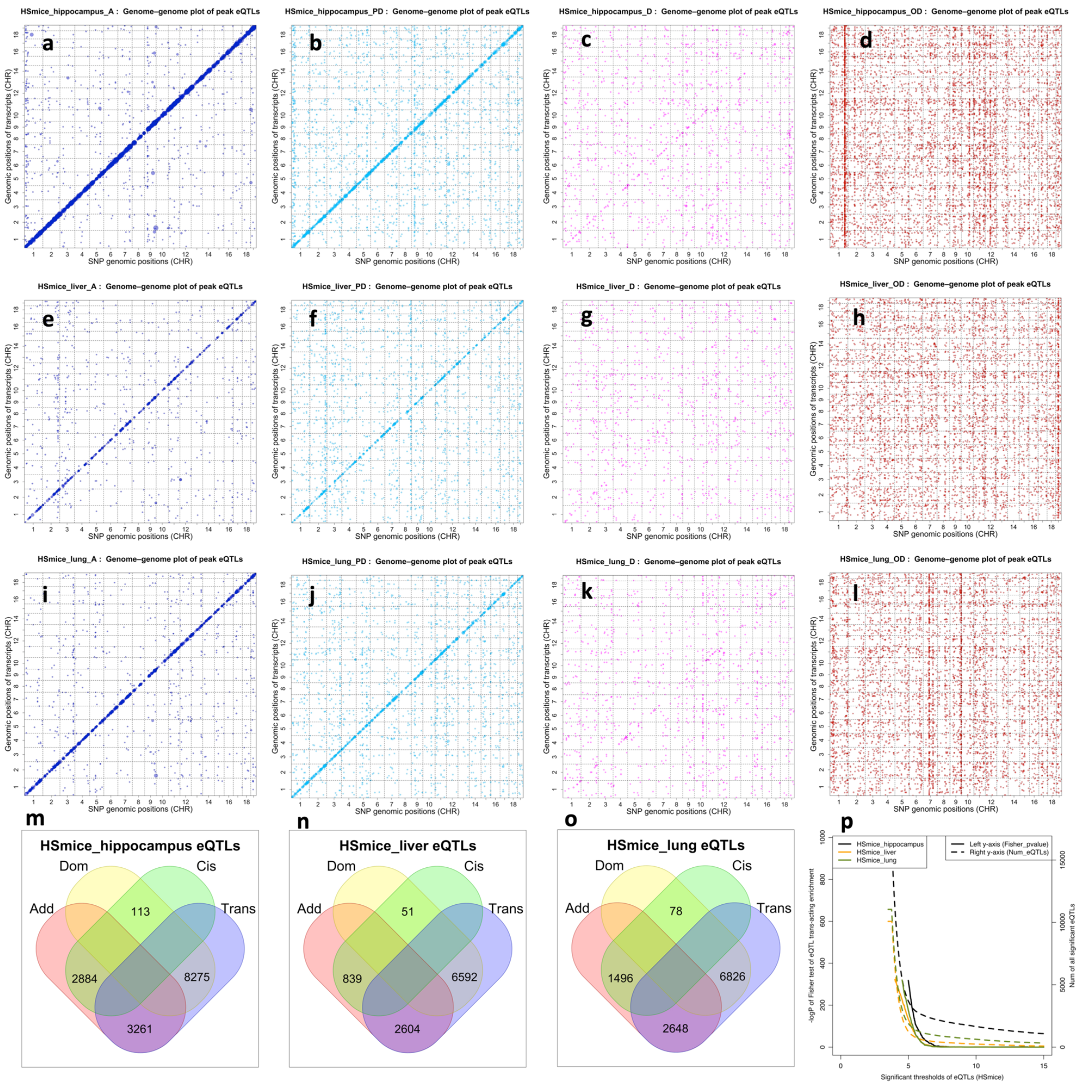


**Fig. S6-3 Trans-acting enrichment among dominant eQTLs of hippocampus, liver and lung tissues in HS mice.** (a-h) eQTL locations of transcript level eQTLs, filtered by dominance type. Each dot represents an eQTL significant at suggestive level (i.e. one false positive expected per transcript). x-axis: eQTL chromosome positions (bp), y-axis: physical gene location. First row (a-d): hippocampus; Second row (e-h): liver; Third row (i-l): lung. The four columns represent dominance types (blue: additive A, sky blue: partial-dominant PD, purple: complete-dominant CD, red: over-dominant OD). (m: hippocampus, n: liver, o: lung) Venn diagrams comparing eQTL cross-classifications, (1) Generalised Additive eQTLs, (A-eQTLs and PD-eQTLs), (2) Generalised Dominance eQTLs, (CD-eQTLs and OD-eQTLs) vs (3) Cis-acting eQTLs, (4) Trans-acting eQTLs. (p) Dominance trans-acting enrichment (left y-axis, solid lines) and the counts of significant eQTLs (right y-axis, dashed lines) under different -log10(P) eQTL significance thresholds (x-axis). Enrichment is quantified by the -log10(P) values of Fisher exact test between dominance (Add/Dom eQTLs) and regulation types (cis/trans-acting) of significant eQTLs within mouse hippocampus (black), liver (orange) and heart (green), respectively.

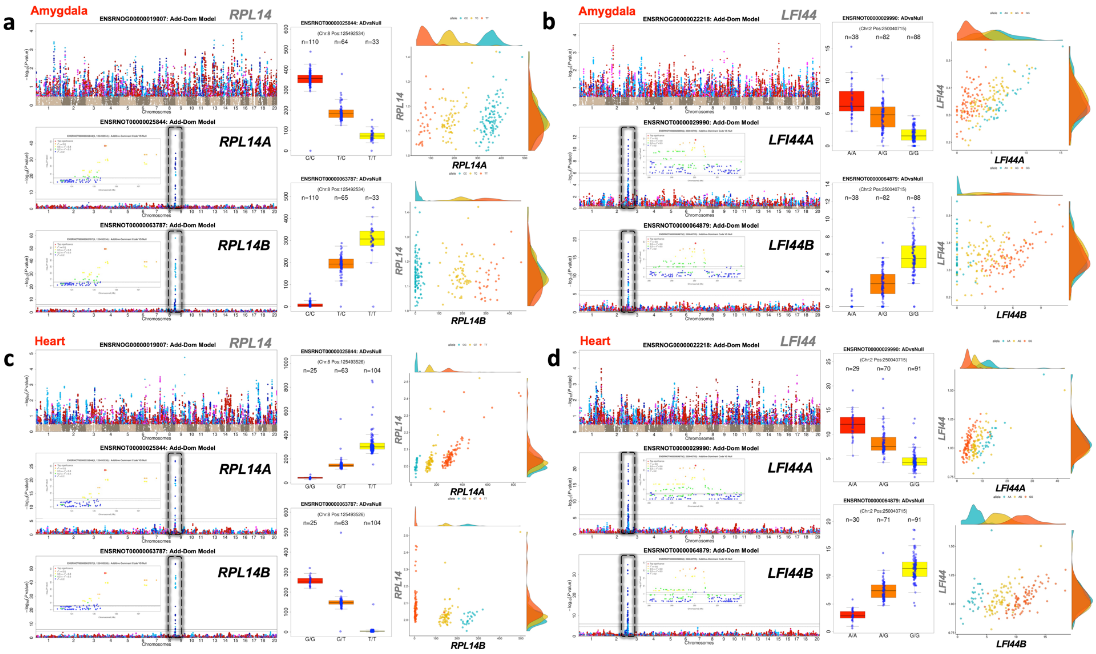


**Fig. S7-1 Transcript-specific antagonistic dominant eQTLs of *RPL14* and *LFI44* in HS rat amygdala and heart.** (a, c) Plots for *RPL14* (a, c) and *LFI44* (b, d) of HS rat amygdala (two upper panels) and heart (two lower panels). On the left part of each panel, the top Manhattan plot is based on their overall gene expression level, showing no genome-wide significant eQTLs, the middle and lower Manhattan plots are based on their two transcripts' expression levels, as well as their corresponding regional plots and phenotypic distribution boxplots of the peak SNPs. Within each Manhattan plot, all the SNPs with $-\log_{10}(P)$ values > 0.5 are coloured by dominance classification. On the right each panel are shown scatter plots of the correlations of expression levels between each transcript with the corresponding overall gene expression. Within each scatter plot, one dot represents one sample, the dot colors indicate the genotype of the peak SNP.

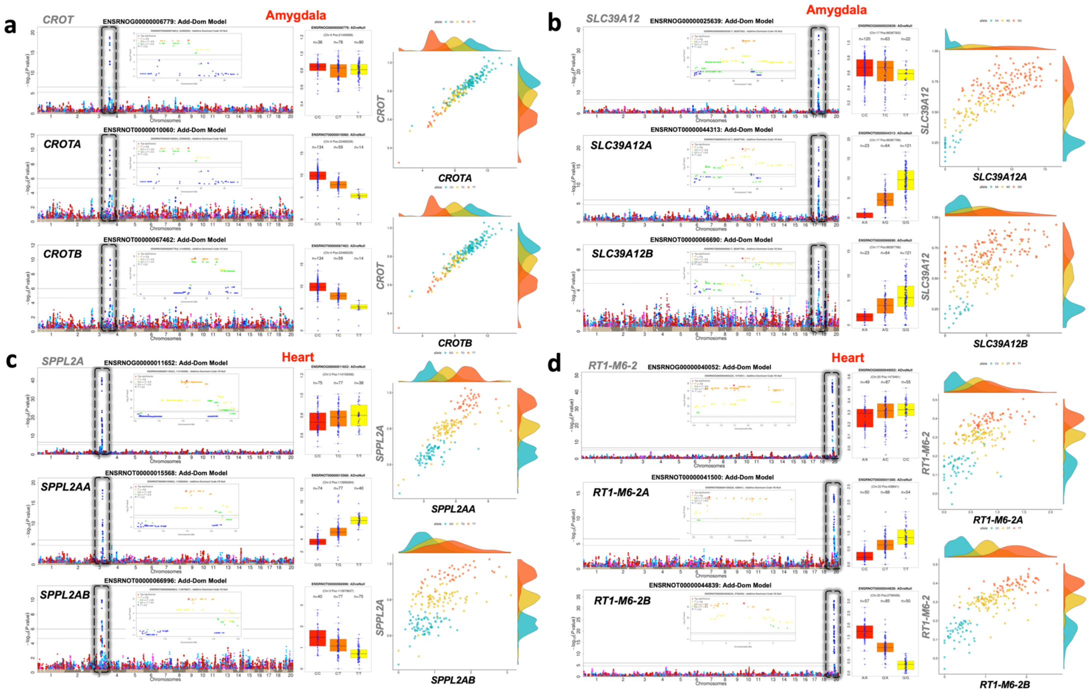


**Fig. S7-2 Transcript-specific synergistic eQTLs in HS rats.** (a-b) Plots of *CROT* (a) and *SLC39A12* (b) of HS rat amygdala. (c-d) Plots of *SPPL2A* (c) and *RT1-M6-2* (d) of HS rat heart. On the left part of each panel, the top Manhattan plot is based on their overall gene expression level, showing no genome-wide significant eQTLs, the middle and lower Manhattan plots are based on their two transcripts' expression levels, as well as their corresponding regional plots and phenotypic distribution boxplots of the peak SNPs. Within each Manhattan plot, all the SNPs with -$\log_{10}(P)$ values > 0.5 are coloured by dominance classification. On the right each panel are shown scatter plots of the correlations of expression levels between each transcript with the corresponding overall gene expression. Within each scatter plot, one dot represents one sample, the dot colors indicate the genotype of the peak SNP.

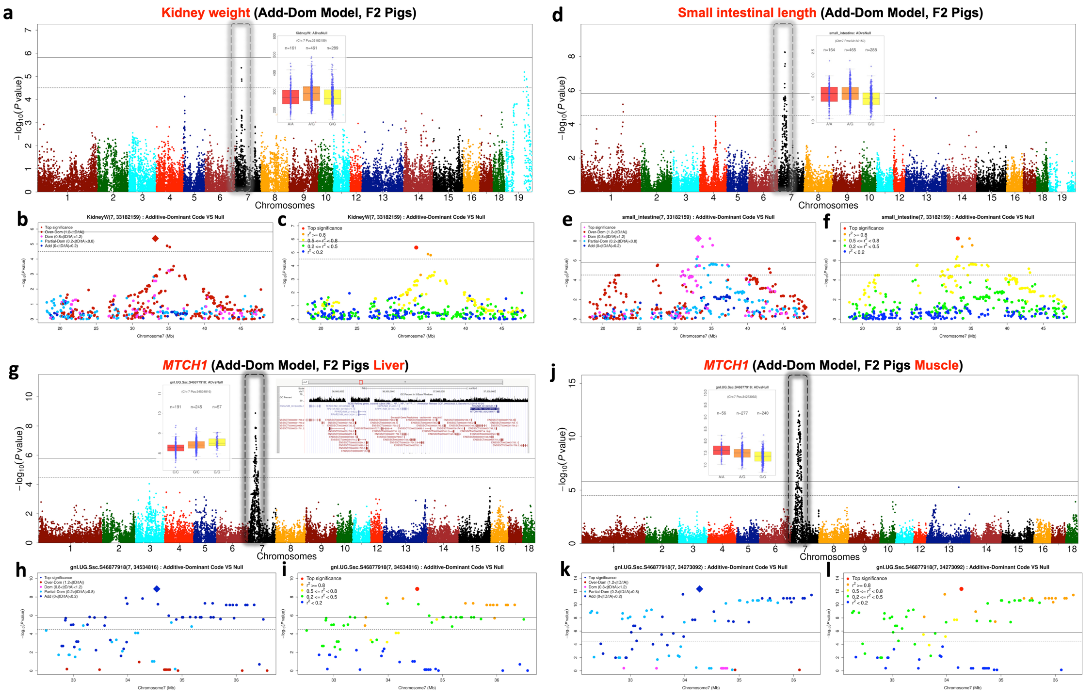


**Fig. S8-1 A pleiotropic over-dominant pig QTL for kidney weight and small intestinal length colocalized with *MTCH1* eQTL in F2 pigs.** Manhattan plots and regional Manhattan plots (chr7: 37Mb-38Mb) (left, blue/red colors represents the QTL dominance types; right, blue/red colors represents the LD ($r^2$) with the peak SNP) are shown based on AD model GWAS for (1) kidney weight GWAS using array SNPs (a-c), (2) small intestinal length using array SNPs (d-f), as well as the *MTCH1* expression level in F2 pigs liver (g-i) and muscle (j-l), respectively. The gene structure plot (g, right upper) from UCSC Genome Browser illustrating *MTCH1* as the nearest gene to the fine-mapped QTL regions of two physical traits (a, d) based on array SNPs.

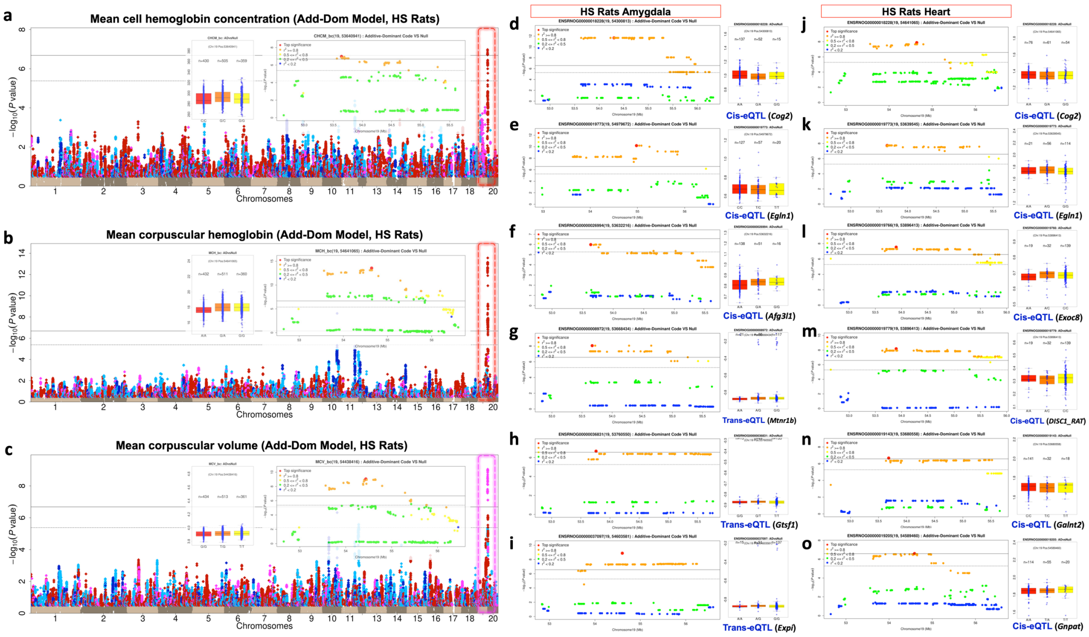


**Fig. S8-2 Dominant QTLs for three rat hematology traits colocalized with cis/trans eQTLs in rat amygdala and heart (chr19: 53 Mb-55 Mb).** (a-c) Manhattan plots, regional Manhattan plots and phenotypic distribution boxplots of the corresponding peak SNPs of three hematology traits in HS rats, (a) mean cell hemoglobin concentration (b) mean corpuscular hemoglobin (c) mean corpuscular volume. Dashed boxes are used to highlight the corresponding QTL regions. (d-i) Regional Manhattan plots and boxplots of colocalized eQTLs of six genes in rat amygdala, including (d-f) three cis-eQTLs of *Cog2* (CD-eQTL), *Egln1* (A-eQTL) and *Afg3l1* (PD-eQTL), and three trans-eQTLs of *Mtnr1b* (CD-eQTL), *Gtsf1* (CD-eQTL) and *Expi* (CD-eQTL). (j-o) Regional Manhattan plots and boxplots of colocalized six cis-eQTLs in rat heart, including *Cog2* (PD-eQTL), *Egln1* (PD-eQTL), *Exoc8* (PD-eQTL), *Disc1_Rat* (PD-eQTL), *Galnt2* (A-eQTL) and *Gnpat* (PD-eQTL). All GWAS based on AD model.

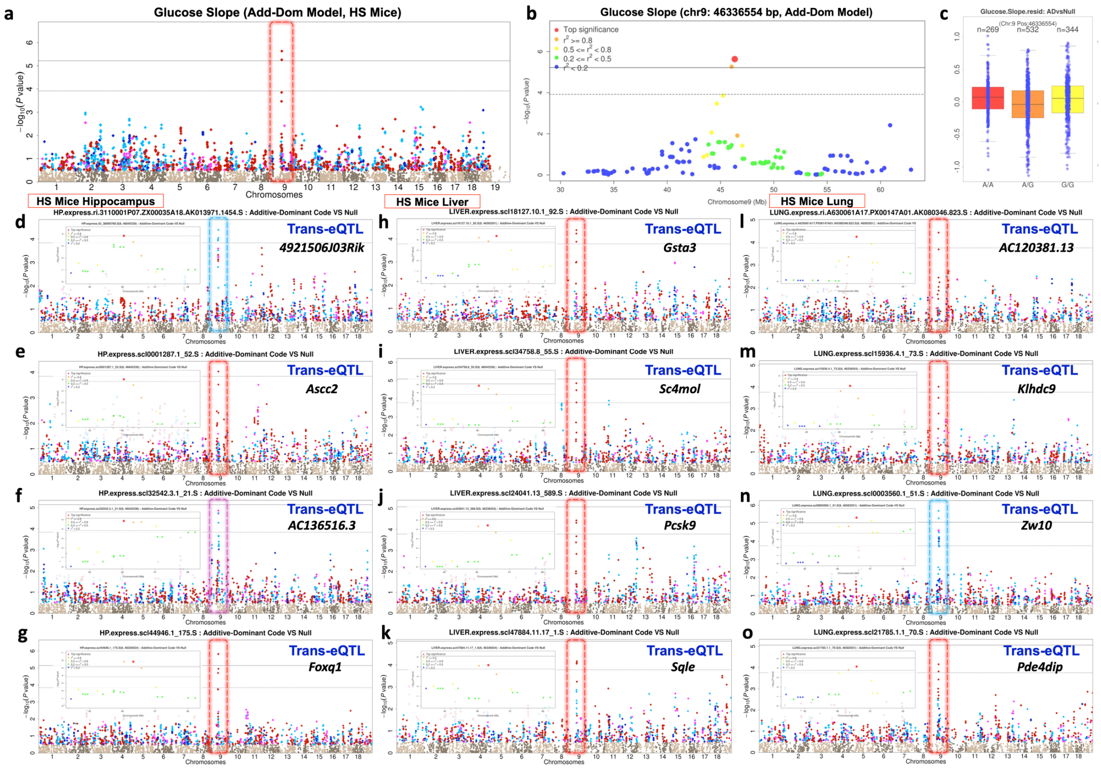


**Fig. S8-3 An over-dominant QTL for mouse glucose slope colocalized with eQTLs in mouse amygdala, liver and lung.** (a-c) Manhattan plots, regional Manhattan plots at (chr9: 45 Mb-47 Mb) and phenotypic distribution boxplots of the corresponding peak SNPs of mean glucose slope in HS mice. (d-g) Manhattan plots and regional plots of four colocalized trans-eQTLs in mouse hippocampus, including *4921506J03Rik* (OD-eQTL), *Ascc2* (OD-eQTL), *AC136516.3* (CD-eQTL) and *Foxq1* (OD-eQTL). (h-k) Manhattan plots and regional plots of four colocalized trans-eQTLs in mouse liver, including *Gsta3* (OD-eQTL), *Sc4mol* (OD-eQTL), *Pcsk9* (OD-eQTL) and *Sqle* (OD-eQTL). (l-o) Manhattan plots and regional plots of four colocalized trans-eQTLs in mouse lung, including *AC120381.13* (OD-eQTL), *Klhdc9* (OD-eQTL), *Zw10* (CD-eQTL) and *Pde4dip* (OD-eQTL). All QTLs based on AD model.
